# Dendritic excitatory-inhibitory balance and branch-specific gating enable selective recall of associative memories

**DOI:** 10.64898/2026.03.27.714865

**Authors:** Severin Berger, Everton J. Agnes

## Abstract

The ability to reconstruct complete memories from fragmented cues is a defining feature of associative memory, which requires neural circuits to store many overlapping and conflicting memory representations while selectively controlling memory accessibility. Attractor dynamics provide the dominant modelling framework for associative memory and have been used to interpret connectivity and behaviour. Yet despite this influence, how attractor-based memory is implemented in biophysical spiking circuits with compartmental dendrites and structured recurrent interactions has remained unresolved, with no unifying theoretical framework for interpreting emerging dendritic imaging and connectomic data. Here we show that local dendritic excitatory-inhibitory balance creates binary-like membrane-potential states that enable associative recall, and that branch-selective inhibition gates access to memory sets stored across dendritic trees, limiting interference between distinct sets. We further show that continuous dendritic dynamics enlarge basins of attraction by integrating partial cues over time, and that a winner-take-all readout circuit implements memory-set selection autonomously, generating interneuron activity that encodes set identity. Our framework predicts branch-specific voltage and calcium signatures that provide a basis for interpreting dendritic measurements in terms of engram organisation, bridging classical associative memory theory with dendritic circuit mechanisms.

---

Memory recall is thought to arise from the reactivation of specific experience-encoding activity patterns in neural circuits^1–7^, often termed *ensembles*^3,4^ or *engrams*^8,9^. How such patterns are retrieved from partial cues—and how circuits select which subset of stored memories is accessible for recall at a given time while suppressing interference from others—remains a central open question. This is particularly challenging in biophysical circuits, where dendrites introduce nonlinear, compartmentalised computations absent from abstract models^10–12^. Attractor models, formalised by Hopfield networks and their variants^13–20^, provide a natural theoretical description of how networks recover stored patterns from noisy inputs and have been widely used to interpret structural connectivity^16,17,21,22^ and behavioural measurements^23,24^. In these models, neurons are typically assigned binary states— active or inactive^14,15^—providing a powerful but abstract representation that leaves open how attractor dynamics are realised in biological circuits.

Existing biophysical implementations of associative memory can form stable cell assemblies, but they typically store only a small number of memory patterns even in large networks^25,26^. Higher capacity has recently been achieved using latent manifolds^27^, but these networks rely on dense all-inhibitory connectivity, limiting their biological realism. Memory recall is also highly dynamic and depends on internal and external background states, often referred to as *context*^28^, which shape memory accessibility—i.e., selective recall, the ability to retrieve specific memories while others stored within the same circuit remain inaccessible. Although contextual gating improves capacity in associative memory models^20^ and supports context-dependent learning in biophysical models^29^, a framework that combines high-capacity associative memory, selective recall, and biophysical realism has remained missing.

Here, we establish a framework that links optimized binary connectivity to biophysical recurrent networks and identify local dendritic excitatory-inhibitory balance as a mechanism for attractor dynamics at the level of dendritic computational subunits. We show that the interplay between linear and nonlinear synaptic currents creates stable dendritic fixed points^30^ that enable binary-like recall in spiking neurons while enlarging basins of attraction through the continuous integration of partial cues. Extending this framework to dendritic trees, we show that branch-selective inhibition gates which dendritic branches can drive somatic spiking, thereby selecting which memory sets are accessible for recall while limiting interference between them. Because this gating requires external control, we further show that a winner-take-all readout network can implement the selection autonomously, stabilising active memory sets and disambiguating highly correlated inputs. The resulting framework predicts branch-specific voltage and calcium signatures that provide a principled basis for linking measurements of dendritic activity^31–33^, compartmentalised plasticity^34,35^, and interneuron population dynamics to the organisation of memory engrams across dendritic trees.

## Results

### EI balance creates dendritic fixed points

We start from a simple biophysical consequence of dendritic synapses: voltage-dependent excitatory and inhibitory currents can naturally create stable membrane-potential fixed points^30,36^ (Fig. 1A,B). For some currents this voltage dependence is set only by the synaptic driving force—i.e., the difference between membrane potential and reversal potential^37^—whereas others exhibit additional voltage-dependent nonlinearities. In particular, N-methyl-D-aspartate (NMDA; Fig. 1Ai, right) and gamma-aminobutyric acid type B activated inward-rectifier potassium currents (GABA_B_/KiR) display nonlinear voltage dependence^30,38–42^ (Fig. 1Aii). The interplay of these nonlinear currents with alpha-amino-3-hydroxy-5-methyl-4-isoxazolepropionic acid (AMPA), gamma-aminobutyric acid type A (GABA_A_), and leak currents generates membrane-potential dynamics with stable and unstable fixed points (Fig. 1B). These fixed points shift systematically with the ratio of excitatory to inhibitory inputs—i.e., presynaptic firing rates weighted by synaptic strengths, rather than the voltage-dependent currents themselves (Fig. 1C). Importantly, a model containing only a spiking somatic compartment cannot access the depolarised fixed point (Fig. 1B, pink lines); explicit dendritic compartments are therefore required for computations that rely on access to both hyperpolarised and depolarised fixed points.

**FIG. 1.**
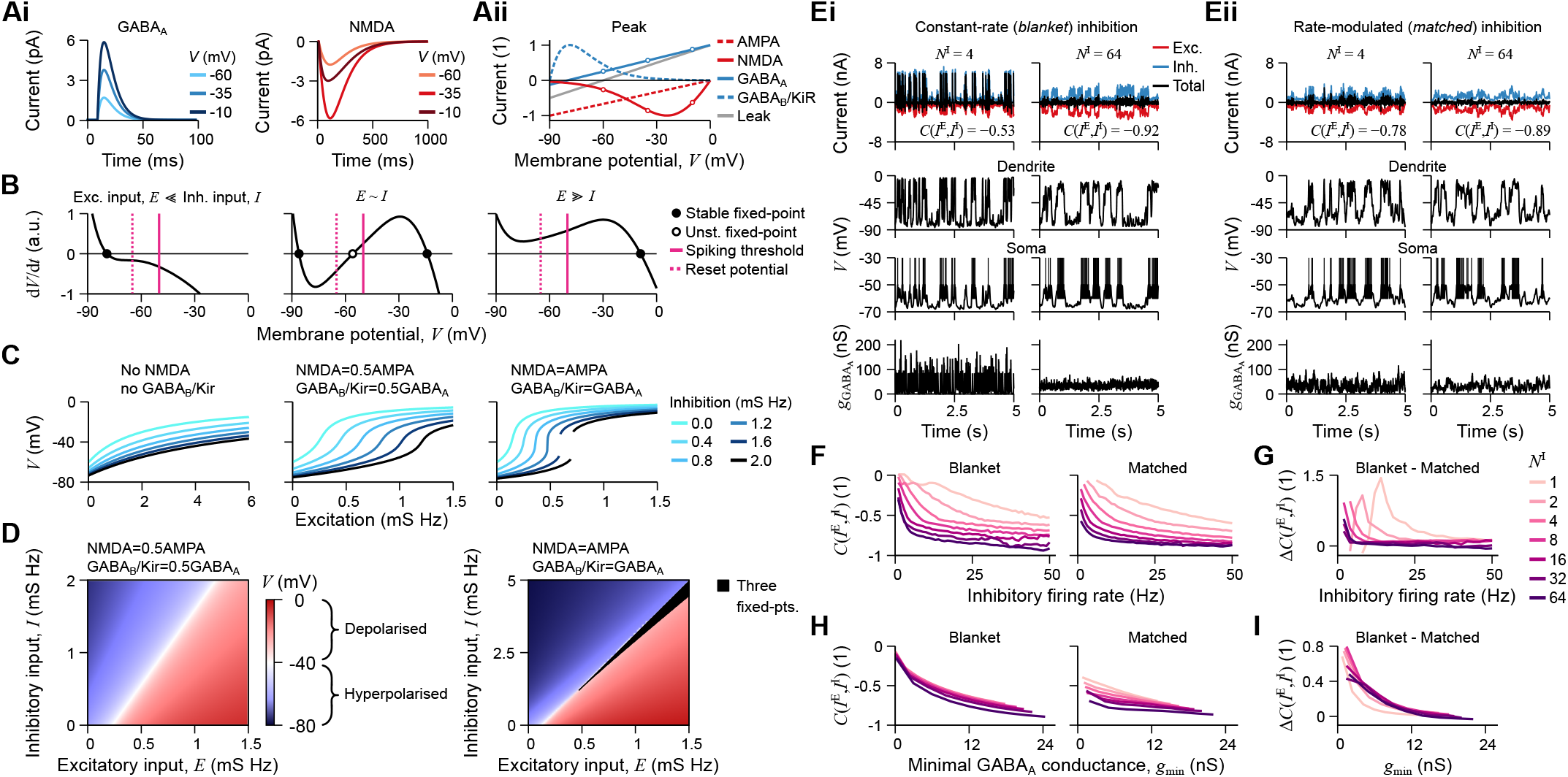
Excitatory-inhibitory balance establishes stable fixed points in dendrites. **A**, Synaptic currents. *Ai*, Example GABA_A_ (left) and NMDA (right) current dynamics for three membrane potentials. *Aii*, Normalised peak synaptic and leak currents as a function of membrane potential, *V*. **B**, Rate of change of membrane potential, d*V/*d*t*, which is proportional to the sum of the currents (cf. panel Aii), as a function of membrane potential, *V*, for three combinations of excitatory and inhibitory input. Zeros of d*V/*d*t* define stable and unstable fixed points. **C**, Stable membrane-potential fixed points as a function of excitatory input for different levels of inhibitory input. The discontinuity marks values of excitatory and inhibitory input for which three fixed points coexist (cf. panel B, middle). **D**, Heat map of membrane-potential fixed points (colour) as a function of excitatory (x-axis) and inhibitory (y-axis) input, yielding a state-space representation of the membrane-potential dynamics. **E**, Excitatory, inhibitory, and total currents (top), dendritic and somatic membrane potentials (middle), and GABA_A_ conductance (bottom) as a function of time. *Ei*, Constant-rate (blanket) inhibition. *Eii*, Rate-modulated (matched) inhibition. The number of inhibitory neurons is denoted by *N*_*I*_. The Pearson correlation between excitatory and inhibitory currents is denoted by *C*(*I*^E^, *I*^I^). **F**, Pearson correlation between excitatory and inhibitory currents, *C*(*I*^E^, *I*^I^), as a function of inhibitory firing rate for different numbers of inhibitory connections, showing that stronger EI balance emerges with larger inhibitory populations or higher firing rates. **G**, Difference in *C*(*I*^E^, *I*^I^) between blanket and matched inhibition as a function of inhibitory firing rate and the number of inhibitory connections. **H, I**, Same as panels F and G, but plotted against the minimum GABA_A_ conductance, *g*_min_.

The relative amplitudes of the nonlinear and linear synaptic currents—i.e., the NMDA-to-AMPA and GABA_B_/KiR-to-GABA_A_ ratios—determine whether membrane-potential dynamics exhibit a single stable fixed point or a bistable regime with two stable fixed points separated by an unstable one (Fig. 1C). Both NMDA and GABA_B_/KiR currents are required for the stable fixed point to acquire an almost binary, step-like dependence on synaptic input (Fig. 1C). We can thus define a two-dimensional state space described by the excitatory and inhibitory input a dendritic branch receives (Fig. 1D). At each point in this state space, the membrane-potential fixed point is defined by the vanishing of the total current, making local excitatory-inhibitory balance the quantity that specifies dendritic state.

### Blanket inhibition balances dendrites

The fixed-point state-space description above (cf. Fig. 1A–D) is defined in terms of temporally averaged excitatory and inhibitory input, raising the question of whether the same balance can be maintained under realistic spiking dynamics. To test this directly, we compared two inhibitory regimes in a single-neuron model: *blanket*^43,44^ and *matched*^45^ inhibition. In both cases, excitatory spikes follow correlated temporal profiles with modulated firing rates^46,47^ (Fig. S1A,B). Under blanket inhibition, inhibitory neurons fire at a constant rate (Fig. S1A), and inhibitory currents become strongly anti-correlated with excitatory currents only when many inhibitory connections converge onto a dendrite (Fig. 1Ei). Under matched inhibition, inhibitory spike times follow the same firing-rate modulation as excitatory spikes (Fig. S1B), so fewer inhibitory connections are sufficient for anticorrelated currents to emerge (Fig. 1Eii). Throughout, we interpret anti-correlated excitatory and inhibitory currents as a signature of EI balance, because anti-correlation indicates that fluctuations in excitation are matched by corresponding changes in inhibition.

Low inhibitory firing rates or too few inhibitory synapses fail to balance excitatory currents in both the blanket and matched conditions. With enough inhibitory neurons or sufficiently high Berger & Agnes (2026) 2 Dendritic balance and memory gating inhibitory firing rates, however, inhibitory currents become anti-correlated with excitatory currents, indicating EI balance (Fig. 1F). Matched inhibitory spike timing provides a clear advantage only when the number of inhibitory neurons is very small and inhibitory firing rates are low (Fig. 1G). The key quantity governing EI balance is the effective minimum inhibitory GABA_A_ conductance (Fig. 1E, bottom; Fig. 1H; Fig. S1C,D; see Methods). Matched inhibitory spike timing is therefore only required when this minimum GABA_A_ conductance is low (Fig. 1I).

A decomposition of the individual synaptic currents (cf. Fig. 1Aii) shows that GABA_A_ can counterbalance NMDA currents in the depolarised regime, whereas GABA_B_/KiR can counterbalance AMPA currents in the hyperpolarised regime. Because the balancing current is given by the product of driving force, non-linear voltage-dependence, and synaptic conductance, the total GABA_A_ conductance on a dendrite (i.e., contribution of all inhibitory synapses), acting on the order of 10 ms per synaptic event^37^, must remain continuously above zero. Maintaining this conductance therefore requires either many convergent inhibitory inputs or sufficiently high inhibitory firing rates. Although inhibitory neurons are fewer in number overall, the high firing rates reported for many inhibitory cell types^48–50^ support the idea that specialised inhibitory physiology is integral to sustaining the minimum conductance required for dendritic EI balance.

### Bi-stable dendrites enable memory recall

The nonlinear dependence of dendritic fixed points on excitatory input creates a sharp transition in dendritic membrane potential (Fig. 2A, top), which in turn produces a steep increase in firing rate over the same input range (Fig. 2A, middle). This input-output relationship approximates the active/inactive dynamics of binary associative memory networks^14,15^ (Fig. 2A, bottom). The state-space description introduced above (cf. Fig. 1D), combined with the somatic spiking threshold, defines a boundary that separates active (depolarised) and inactive (hyperpolarised) dendritic states as a function of excitatory and inhibitory input (Fig. 2B), with each state corresponding to a distinct postsynaptic firing rate (Fig. S2). This allows each neuron to be approximated as either inactive, producing no output spikes, or active, firing at a defined rate above an inhibition-dependent input threshold. On this basis, we derive a binary-to-biophysical mapping that embeds associative memory dynamics in biophysical spiking networks (see Methods for details).

**FIG. 2.**
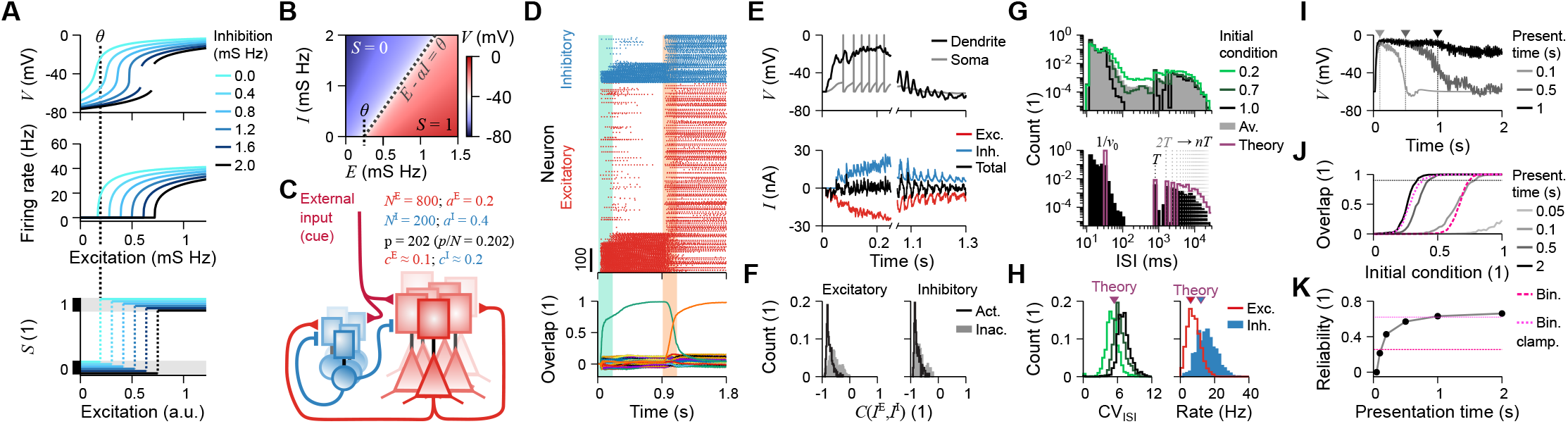
Binary-to-biophysical mapping enables associative recall through dendritic EI balance. **A**, Membrane-potential fixed points, *V* (top; cf. Fig. 1C), the resulting firing rate (middle), and the corresponding binary state (bottom) as a function of excitatory input for different levels of inhibition. **B**, State-space of dendritic membrane potential (cf. Fig. 1D). The dashed line separates hyperpolarised (*S* = 0) and depolarised (*S* = 1) states. **C**, Schematic of the recurrent excitatory-inhibitory network. Key parameters shown: *N*, number of neurons; *a*, activity level; *c*, connection density; *p*, number of stored patterns. Superscripts E and I denote excitatory and inhibitory populations. **D**, Spike times of excitatory (red) and inhibitory (blue) neurons (top) and memory overlap (bottom) as a function of time. Shaded areas indicate periods of external stimulation. **E**, Examples of membrane potential (top) and currents (bottom) during active (left) and inactive (right) periods of a neuron during recall of two memory patterns (cf. panel D). **F**, Histograms of *C*(*I*^E^, *I*^I^) for excitatory (left) and inhibitory (right) neurons, separated into active (black) and inactive (grey) populations. **G**, Distribution of interspike intervals (ISIs) for different initial condition sizes, defined as the fraction of the memory pattern that is externally activated. Key parameters: *ν*_0_, firing-rate during active recall; *T*, period of external activation; *nT*, interval spanning *n* activation periods. **H**, Distribution of the coefficient of variation of interspike intervals, CV_ISI_ (left), and neuronal firing rates (right). **I**, Examples of dendritic membrane-potential trajectories for different presentation times. **J**, Overlap between the final network state and the target memory pattern as a function of initial condition size (cf. panel G). **K**, Reliability of memory recall, defined as one minus the smallest initial condition size for which the average overlap exceeds 0.9 (from panel J), as a function of presentation time. Higher reliability indicates that smaller fractions of a memory pattern are sufficient to trigger full recall. Pink lines show the equivalent binary network without (dashed) and with (dotted) clamped initial-condition units.

To test this binary-to-biophysical mapping, we trained a binary network to store memory patterns using a modified perceptron learning rule (modPLR), based on previous work^17,51,52^, and mapped the resulting connectivity onto our biophysical spiking network (Fig. 2C; see Methods). The modPLR allows us to maximise memory capacity while constraining synaptic density to match biological observations (Figs. S3 and S4A). The mapped biophysical spiking network exhibits robust attractor dynamics and reliably recalls memories from strongly corrupted inputs (Fig. 2D). To recall a given memory, a subset of neurons corresponding to a corrupted version of that memory is driven for a defined period by external excitatory input in the form of Poisson spikes (shaded green and orange backgrounds in Fig. 2D). When a neuron belongs to the recalled memory, it becomes active and receives strong but balanced currents (Fig. 2E, left). The same neuron, when it does not belong to the recalled memory, remains inactive, with hyperpolarised dendritic and somatic membrane potentials and weak but balanced currents (Fig. 2E, right). As in the single-neuron analysis (cf. Fig. 1E), neurons in the recurrent network receive anti-correlated excitatory and inhibitory currents (Fig. 2F), consistent with a state of EI balance. When multiple patterns are recalled in sequence by successive external activations, interspike intervals (ISIs) show a bimodal distribution (Fig. 2G), together with high coefficients of variation and low firing rates (Fig. 2H), consistent with activity reported in hippocampal^53–55^ and cortical^56^ neurons.

Membrane-potential dynamics unfold over tens of milliseconds, allowing dendritic activation to integrate and persist over time (Fig. 2I). This differs markedly from binary networks, in which neurons integrate their inputs and update instantaneously at each time step, and from single-compartment spiking neurons, which reset after each spike and cannot access the depolarised fixed point (cf. Fig. 1B, pink lines). The longer external stimulation keeps a partial memory pattern active, the more likely it is that additional neurons belonging to that memory are recruited. As a result, even small externally activated fractions of a memory pattern, which we define as the initial condition size, can trigger full recall if the stimulation lasts long enough (Figs. 2J and S4B). Across all initial condition sizes, recall performance increases with presentation time: 100 ms matches binary-network performance, whereas approximately 1 s matches the performance of binary networks with clamped units (Figs. 2K and S4B).

These results establish that optimized binary connectivity can be mapped onto biophysical spiking networks through stable dendritic membrane-potential fixed points that arise directly from synaptic current dynamics. This mapping remains robust across a broad range of network parameters (Figs. S5 and S6; see Supplementary Text). Recall remains reliable in the biophysical model whenever the stability of memory patterns in the binary network exceeds a threshold (Fig. S5F), although performance decreases as the number of stored patterns increases (Fig. S5G). Beyond fixed-point recall, the same mapping also supports autonomous sequential recall when connectivity is optimized to embed transitions between memory patterns^17^ (Fig. S7). This establishes a biophysical realisation of classical attractor-memory theory and links dendritic integration to collective network dynamics.

### Inhibition gates selective recall

Distributing memory storage across dendritic branches creates the possibility of selective recall. Adding a second dendritic compartment (Fig. 3A) expands the computational capacity of the network by enabling additional memories to be stored and selectively accessed^20^. Assigning an independent weight matrix to this second dendrite separates storage into two memory sets (Fig. 3Bi). Without further regulation, however, these two sets interfere strongly and recall fails (Fig. 3C).

**FIG. 3.**
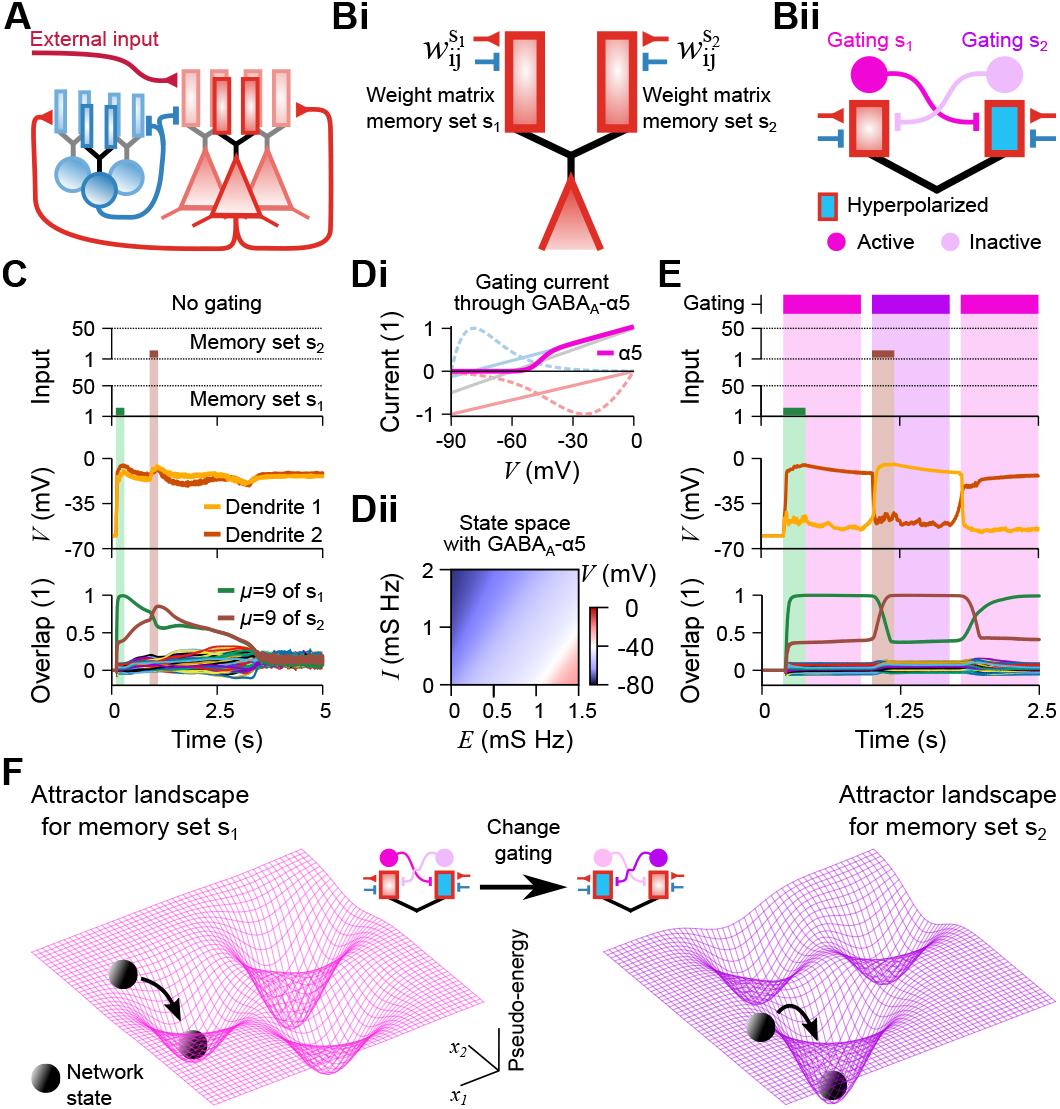
Targeted dendritic inhibition switches which memory set is available for recall. **A**, Schematic of the simulated network with a second dendrite (extension of Fig. 2C). **B**, Memory storage and inhibitory gating. *Bi*, Schematic of a neuron with two distinct connectivity matrices, one per dendrite, 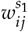 and 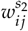, storing two sets of memory patterns, *s*_1_ and *s*_2_. *Bii*, Schematic of inhibitory gating. External inhibitory connections target opposite dendrites, so that recalling memories stored in one dendrite requires shunting the other dendrite across the network. **C**, Temporal evolution of the input (top), dendritic membrane potential (middle), and network overlap with the stored memories (bottom) without inhibitory gating. Shaded areas indicate periods of external stimulation (cf. Fig. 2D). **D**, Gating through GABA_A_-*α*5 synapses. *Di*, Same as Fig. 1Aii, but with an added GABA_A_-*α*5 peak current. *Dii*, Same as Fig. 1D, but with constant GABA_A_-*α*5 input. **E**, Same as panel C, but with active inhibitory gating. Narrow shaded bars indicate periods of external stimulation (as in panel C), and broad coloured backgrounds indicate which memory set is accessible for recall: pink when *s*_1_ is accessible (dendrite 2 shunted) and purple when *s*_2_ is accessible (dendrite 1 shunted). **F**, Schematic showing that inhibitory gating switches which attractor landscape is accessible. *x*_1_ and *x*_2_ denote hypothetical reduced neuronal dimensions, and the vertical axis denotes the pseudo-energy of the system.

To resolve this interference, we introduced a second class of inhibitory connections that gates dendrites by shunting one branch, leaving only the other free to integrate excitatory input (Fig. 3Bii). For this gating mechanism, we modelled nitric-oxide-synthase-expressing (NOS) neurogliaform cells^57^ with an activation non-linearity that effectively shunts the targeted dendrite (Fig. 3D). By controlling dendritic accessibility over time, this externally applied gating enables selective recall of two independent memory sets without interference (Fig. 3E).

Inhibitory gating has two main consequences for network performance. First, distributing a fixed total number of synaptic connections across multiple dendrites slightly reduces overall memory capacity because of increased synaptic sparsity^17^ (Fig. S8A), but increases the stability of individual stored memories (Fig. S8B,C). Second, gating renders the attractor structure of the network dynamic^20^. Switching the inhibitory state changes which memory set defines the stable basins of attraction available for recall (Fig. 3F; see Supplementary Text).

To extend selective recall across multiple memory sets, memories can be distributed across a complex dendritic tree (Fig. 4A), provided that active dendritic amplification counteracts passive attenuation and preserves synaptic democracy^58^ (Fig. S9). In our framework, memory-encoding synapses are confined to terminal dendritic branches (leaves), and recalling a given memory set requires branch-specific shunting inhibition to select the corresponding leaf (Fig. 4B). As additional memory sets are stored, more extensive inhibitory gating is required to silence inactive branches (cf. Fig. 3). We therefore considered three gating strategies that select a single active leaf (Fig. 4C): inhibiting all off-pathway dendrites (“All”), inhibiting only the inactive leaves (“Memory-set-specific”), or inhibiting only the branches that share branching points with the active path (“Optimal”). The last strategy is optimal in the sense that it minimises the number of gated branches required to isolate a memory (Fig. 4D). Both the “All” and “Optimal” strategies require inhibitory synapses to be distributed across the dendritic tree, matching structural observations in hippocampal pyramidal neurons^59^.

**FIG. 4.**
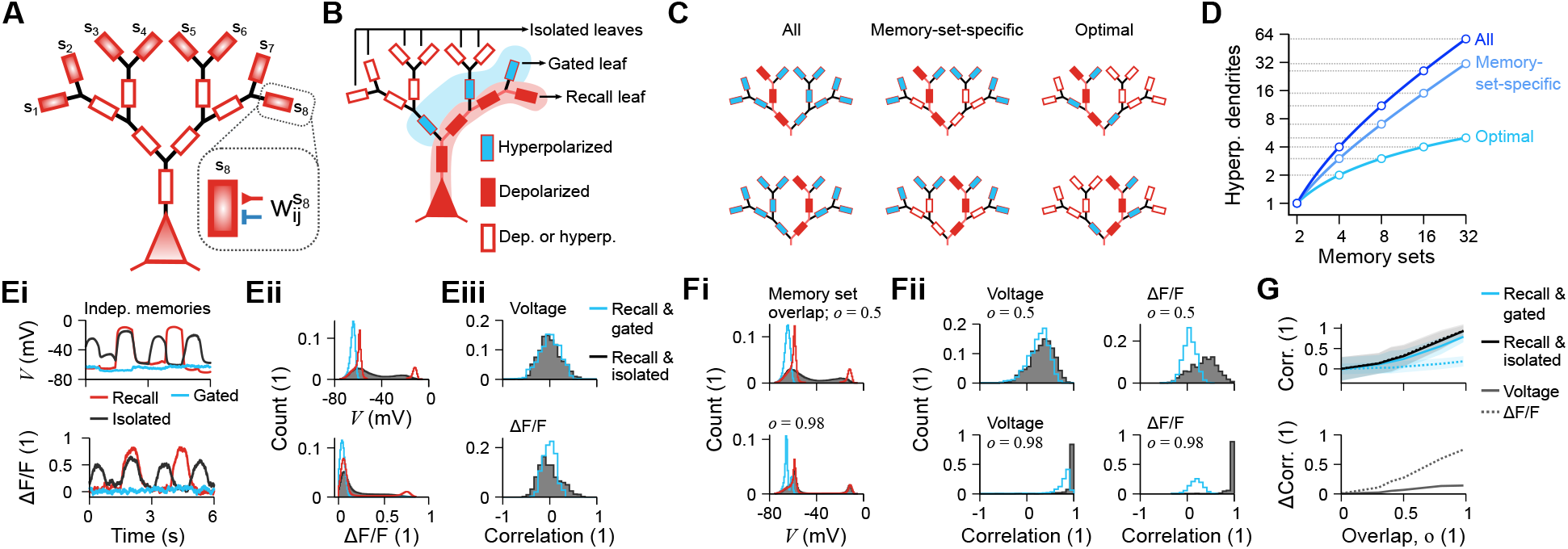
Complex dendritic trees predict branch-specific signatures of inhibitory gating. **A**, Complex dendritic tree in which each branch bifurcates into two branches. Each leaf dendrite receives a distinct weight matrix, for example 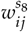. **B**, Neuron with one recall leaf dendrite driving somatic output, one gated leaf, and isolated leaves that may be either depolarised or hyperpolarised. **C**, Three inhibitory gating strategies that select a single recall leaf dendrite (to ensure somatic access): gating all dendrites (All), only leaf dendrites (Memory-set-specific), or only branches sharing a branching point with the active path (Optimal). **D**, Number of hyperpolarised dendrites as a function of the number of memory sets for the strategies in **C. E**, Activity comparison for independent memories across three leaf dendrites: recall, gated, and isolated. *Ei*, Temporal evolution of membrane potential (top) and the corresponding fluorescence signal (bottom). *Eii*, Distributions of membrane potential (top) and fluorescence signal (bottom). *Eiii*, Pearson correlation distributions for membrane potential (left) and fluorescence (right) between the recall dendrite and gated or isolated dendrites. **F**, Activity comparison for overlapping memory sets. *Fi*, Distributions of membrane potential for memory-set overlap *o* = 0.5 (top) and *o* = 0.98 (bottom). *Fii*, Pearson correlation distributions for membrane potential (left) and fluorescence (right) for *o* = 0.5 (top) and *o* = 0.98 (bottom). **G**, Mean Pearson correlation between the recall dendrite and gated or isolated dendrites as a function of overlap (top), and the difference between recall-gated and recall-isolated pairs as a function of overlap (bottom).

In our framework, when a given memory set is recalled, only the leaf dendrite storing that set influences somatic output—the *recall* leaf dendrite (Fig. 4B). Under the optimal gating strategy (Fig. 4C), only the adjacent leaf is explicitly gated off—the *gated* leaf dendrite. As a result, other non-recall leaves can still become depolarised by network activity without affecting somatic spiking—the *isolated* leaf dendrites—because intermediate trunk dendrites connecting these non-recall leaves to the soma are shunted. These compartmentalised dynamics could be tested experimentally using targeted measurements of dendritic membrane potential^31,33^ or calcium fluorescence imaging (Δ*F/F*)^32,33^, which should reveal localised transients in isolated leaves driven by concurrent network activity (Fig. 4Ei).

We predicted the statistical signatures of inhibitory gating by comparing the recall, gated, and isolated leaves (cf. Fig. 4B) in networks storing either independent (Fig. 4E) or overlapping (Fig. 4F) memory sets. In both cases, the membrane potential of the recall leaf follows a bimodal distribution whose depolarised mode reflects the sparsity of the memory patterns (for example, depolarised for roughly 20% of the time when activity is 20% sparse) (Fig. 4Eii, Fi; red). By contrast, the adjacent gated leaf remains hyperpolarised (Fig. 4Eii, Fi; blue), whereas the isolated leaf shows a broad and fluctuating voltage distribution driven by general network activity (Fig. 4Eii, Fi; black). Simulated calcium fluorescence broadly mirrors these voltage distributions, but differs from membrane voltage in its cross-branch correlations (Fig. 4Eiii, Fii). In particular, small subthreshold fluctuations in the shunted leaf remain correlated with the recall dendrite at the level of membrane voltage, whereas calcium fluorescence acts as a slower nonlinear filter that is dominated by large depolarisations (Fig. 4Ei). As a result, fluorescence correlations between recall and gated leaves fall to near zero. By filtering out shared subthreshold fluctuations, dendritic fluorescence provides a clearer and higher-contrast signal than membrane voltage for identifying the degree of structural overlap between stored memory sets (Fig. 4G). Distinct architectures of overlap across dendritic memory sets also generate different predictions for synapse-per-pair statistics, providing an additional structural signature of how memories are distributed across dendritic trees (Fig. S10; Supplementary Text).

These results identify branch-specific activity and correlation patterns as experimentally accessible signatures of how memory sets are distributed across dendritic trees. For this gating mechanism to operate without external control, the circuit itself must determine which dendrite is active.

### Readout networks control memory access

To make dendritic gating an intrinsic circuit mechanism rather than an externally imposed control, we constructed a closed-loop system of three interconnected networks (Fig. 5A). This architecture converts selective recall into an autonomous circuit computation by coupling the core memory network to dedicated readout and gating circuits. The core memory network is configured as above, with recurrent excitatory connectivity derived using our modified perceptron learning rule. Its excitatory neurons project to a readout network in which local inhibitory neurons implement winner-take-all competition so that only one memory pattern is represented at a time (Fig. 5A,B). To mirror the microcircuitry of hippocampal area CA1^60^, we omitted recurrent excitatory-to-excitatory connections within the readout network, although this constraint can be relaxed to model, e.g., CA3 laminar organisation^61^ or cortical circuitry^62^. Both memory-to-readout and recurrent readout connections were derived using the same modified perceptron learning rule, yielding a readout circuit with only excitatory-to-inhibitory and inhibitory-to-excitatory recurrent connections—a simple lateral-inhibition architecture. For simplicity, we assigned non-overlapping representations to readout neurons (Fig. 5B), although the same architecture is compatible with distributed coding. Finally, gating inhibitory neurons receive excitation from the readout network and return shunting inhibition to the dendrites of the memory network, thereby closing the loop (Fig. 5B; cf. Fig. 4C).

**FIG. 5.**
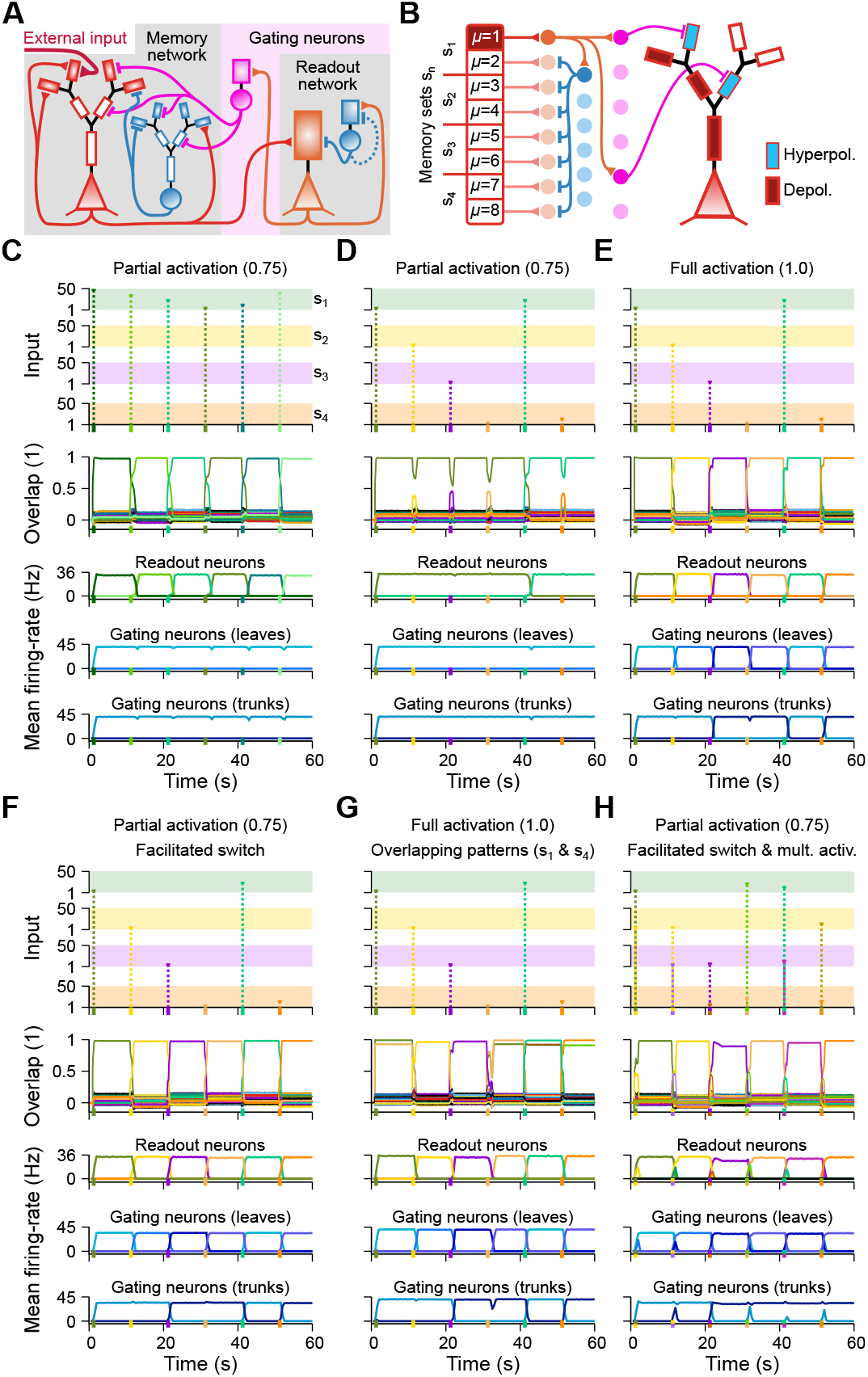
An autonomous readout network gates dendrites to prevent memory interference. **A**, Schematic of the coupled-network architecture, comprising a memory network, a readout network, and gating neurons. **B**, Schematic of the readout and gating connectivity. Readout neurons representing the recalled memory activate gating neurons, which in turn shunt selected dendrites in the memory network. **C**, Temporal evolution of external input, memory overlap, and the mean firing rates of readout and gating neurons during sequential partial activation of six memories from set *s*_1_. Each memory is activated by external stimulation of 75% of its neurons during a 500 ms period indicated by the dashed line. **D**, Same as panel C, but for partial activation of memories from multiple sets. Partial activation of memories from a different set does not switch the active gating state. **E**, Same as panel D, but with full external activation of each memory. Full activation is sufficient to switch the active memory set and the corresponding gating state. **F**, Same as panel D, but with reduced recurrent connectivity in the readout network. This facilitates switching between memory sets under partial activation. **G**, Same as panel E, but for highly overlapping memory patterns from sets *s*_1_ and *s*_4_. The readout network selects a single memory representation despite overlap in the memory network. **H**, Same as panel F, but with simultaneous partial activation of two memories from different sets. The network resolves the competition by selecting a single readout and gating state.

We first stored independent memory patterns and tested their recall by externally activating the memory network (Figs. 5C–E and S11A–C). Starting from a silent state, partial activation is sufficient to trigger recall of an initial memory, as shown by both memory overlap and readout activity (Fig. 5C). The readout neurons then recruit a subset of gating inhibitory neurons (Fig. 5C), which shunt the corresponding dendrites in the memory network (Fig. S11A). Subsequent partial activation of other memories from the same set also induces full recall (Fig. 5C). Once a memory set has been established in this way, partial activation of a memory from a different set is insufficient to overcome the current gating state (Fig. 5D). Instead, full activation of memories from distinct sets is required to switch the gating state and enable recall of the corresponding memories (Fig. 5E). Reducing the strength of recurrent excitatory-to-inhibitory and inhibitory-to-excitatory connections in the readout network allows partial activation to switch memory sets fully (Figs. 5F and S11D). Thus, autonomous recruitment of gating neurons by the readout network stabilises the active memory set against interference from competing sets.

Beyond protecting active memory sets, the winner-take-all readout also resolves ambiguity between highly overlapping patterns. When two memory sets contain strongly overlapping memories (*o* = 0.9), activity in the memory network becomes highly redundant, as reflected by the overlap measure (Figs. 5G and S11E). Despite this redundancy, the readout network selects a single memory representation (Fig. 5G), which in turn drives the corresponding gating feedback. When memories from different sets are instead activated simultaneously, competition in the readout network ensures that only one memory is fully recalled and represented (Figs. 5H and S11F). The readout network thus transforms redundant memory-network activity into a unique pattern selection, even when the underlying representations are nearly indistinguishable.

Notably, the activity of gating neurons encodes memory-set identity rather than individual memory patterns (Fig. 5C–H, bottom), providing a set-level signal that may not be directly available from the memory network itself. Our framework therefore predicts that the dendrite-targeting interneuron population that controls memory-set selectivity—such as NOS neurogliaform cells^38,57^— should exhibit sustained activity that is invariant to which memory is recalled within a set, but switches sharply when a different memory set becomes accessible. This encoding does not need to be as localised as in our implementation; a distributed representation across gating interneurons would suffice, provided it consistently distinguishes between memory sets. Together, these results establish autonomous readout-gating as a circuit mechanism for controlling memory accessibility, protecting active engrams from interference, and disambiguating overlapping inputs.

## Discussion

Our work establishes a biophysical route by which classical associative-memory dynamics can be implemented in spiking neural circuits. We show that local dendritic membrane-potential fixed points can support attractor recall even under a “blanket of inhibition”, where inhibitory spikes do not need to precisely track fast excitatory fluctuations. By providing a mathematical bridge between classical attractor-memory theory^13,14,16–18^ and biophysical circuit mechanisms, our framework reframes dendritic properties—often viewed as complications for network computation—as the substrate that enables both high-capacity recall and selective memory access through local EI balance and branch-specific gating.

EI balance is a cornerstone of many network models and is also an actively regulated property of biological circuits^63,64^, but its mechanistic implementation varies widely. Previous frameworks have relied on macroscopic statistical cancellation to generate chaotic asynchronous states^65^, precise microscopic spike tracking to minimise representational errors^66^, or finely tuned feedback to stabilise strongly recurrent networks—either globally^67^ or through state-dependent supralinear scaling^68^. Recent work has also shown that precise balance can shape the geometry and dynamics of memory representations in recurrent networks^69^, but this too operates at the somatic level. In contrast to these network-level or somatic formulations, EI balance in our framework emerges locally as dendritic membrane-potential fixed points. These fixed points arise from the interaction between distinct synaptic currents and their voltage-dependent driving forces, showing how local biophysics can stabilise memory computations without requiring instantaneous spike-by-spike cancellation.

The computational roles required by our framework— dendritic inhibition, gating, and blanket inhibition—map onto distinct interneuron classes with known connectivity and physiology^48,60,61,70,71^. In our framework, memory-related inhibition targets leaf dendrites, consistent with the connectivity of somatostatin-expressing (SOM) interneurons^48,60^. Controlling memory accessibility further requires a second class of gating interneurons, for which NOS neurogliaform cells provide one plausible candidate implementation^38,57,59^. More generally, the interplay between these interneuron classes suggests how dendritic inhibition and disinhibition could be coordinated across cell types^72^. Because direct SOM-to-SOM connectivity is sparse, this coordination could also involve vasoactive intestinal peptide (VIP) neurons, which provide targeted disinhibition of SOM networks^60,73^. Fast-spiking parvalbumin (PV) neurons could, in turn, contribute to the blanket of inhibition required by the model^44,70^. Different circuit motifs could in principle implement the same computational roles, so the mappings proposed here should be viewed as candidate solutions rather than unique biological assignments. In particular, our results predict that the population of dendrite-targeting gating interneurons should carry a sustained signal encoding memory-set identity that is invariant to which individual memory is recalled. Our framework therefore generates testable hypotheses about how specialised interneuron populations may support dendritic memory gating in biological circuits.

Memory accessibility is highly dynamic, with external cues recalling different memories depending on context^28,74,75^. Recent work further suggests that hippocampal representations are tuned to the computational demands of context-rich memory tasks and that dendritic modulation can support such context-dependent computations^76,77^. In our framework, memory sets stored in separate dendritic leaves provide a natural substrate for such functional contexts. In hippocampal circuits, this contextual gating could in principle be influenced by feedforward input from the dentate gyrus (DG)^78,79^ or by feedback from CA1 and cortex^59,74,80,81^, and the coexistence of functionally distinct excitatory populations with specific inhibitory interactions within CA3 itself^61^ suggests that memory and readout computations do not need to reside in separate areas. Disruption of this gating mechanism suggests links to pathological memory states. For example, the artificial recovery of “lost” memories in Alzheimer’s models^82^ is consistent with normally inaccessible memories becoming retrievable when inhibitory gating is altered, and disrupted hippocampal bursting in Alzheimer’s-related pathology^83^ could further destabilise selective recall. Conversely, weakened gating could allow inappropriate memory sets to become accessible in safe contexts—as observed in post-traumatic stress disorder (PTSD), where traumatic memories intrude despite contextual safety^84^. More broadly, conditions that shift the GABA_A_ reversal potential—such as disrupted chloride regulation in Down syndrome^85^—may compromise the dendritic EI balance required for stable fixed points, and thereby also impair both recall and gating.

Our model uses mathematically optimised connectivity matrices as a benchmark for binary associative memory networks^16,17^. How such structured connectivity is learned *in vivo* remains unresolved. One possibility is that it emerges through compartmentalised learning^35,86^, driven by dendrite-specific plasticity rules^29,47,87–91^ and shaped by distinct inhibitory plasticity mechanisms^46,63,92^. Epigenetic regulation of neuronal excitability^93,94^ may further influence which neurons are allocated to specific engrams, thereby determining the structure of the memory patterns that our framework takes as given.

Our work isolates the minimal ingredients required to explain a specific problem: how selective recall of memory attractors can be implemented in biophysical spiking circuits. Optimized connectivity isolates the mapping between classical associative-memory solutions and biophysical networks^16,17^; branch-specific shunting inhibition isolates the access-control problem^20,57^; and active dendritic amplification isolates the conditions under which selective recall can scale across dendritic trees^58^. From this perspective, the autonomous readout circuit should be interpreted as a biologically motivated proof of principle rather than a unique circuit realisation. Although we use CA3 and CA1 as motivating examples for the memory and readout networks, respectively, the same computational architecture could be realised within a single area—for example, across layers of CA3 or cortex^48,62^, or among functionally distinct neuron types within the same region^5,61^. Similarly, the experimental links developed here are intended not as fits to a particular dataset, but as structural and functional signatures that can distinguish this mechanism from alternative implementations^32,34^. For instance, the active dendritic amplification required for selective recall across complex dendritic trees is known to be suppressed during general anaesthesia^95^, illustrating how specific biophysical conditions can gate the mechanism as a whole. An important next step will be to determine how closely biological circuits approximate these requirements and how such organisation could emerge through development^5,96^ and learning^35,47^, particularly given recent evidence that hippocampal neurons can encode large numbers of sparse, distinct memory representations^6^.

Our work establishes dendritic balance and branch-selective gating as sufficient circuit mechanisms for selective attractor recall in biophysical spiking networks. By grounding associative-memory dynamics in dendritic physiology, our framework transforms the dendritic tree from a passive recipient of synaptic input into an active organiser of memory accessibility—with branch-specific voltage and calcium signatures that make this organisation experimentally visible. As tools for measuring and manipulating dendritic activity in vivo continue to advance, the framework developed here provides a theoretical foundation for reading out how memories are distributed, selected, and protected across the dendritic trees of single neurons.

## Acknowledgements

We thank Josef Bischofberger for discussions on the role of cell-type-specific interneurons in dendritic gating. This research was funded by the Swiss National Science Foundation (SNSF) [10005175]. For the purpose of Open Access, a CC BY public copyright licence is applied to any Author Accepted Manuscript (AAM) version arising from this submission.

## Software and code availability

Code will be made available upon publication.

## Methods

### Biophysical model

#### Dendritic compartment

The membrane potential, *V*_*i*_(*t*), of dendritic compartment *i* in a neuron (*i* = 1, …, *N*^D^, where *N*^D^ is the total number of dendritic compartments) evolves according to

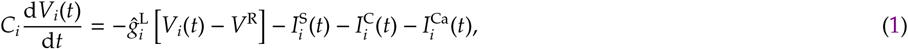

where *C*_*i*_ = *C*Δ*A*_*i*_ and 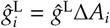 are the total capacitance and leak conductance of compartment *i*, obtained by multiplying the capacitance per unit area, *C*, and leak conductance per unit area, *ĝ*^L^, by the branch area, Δ*A*_*i*_. This choice gives all dendritic compartments the same membrane time constant, *τ*^M^ = *C/ĝ*^L^. Throughout, hatted conductances denote absolute conductances (for example, in nS), whereas unhatted conductances denote conductances normalized by the compartment-specific leak conductance, 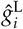. In Eq. 1, *V*^R^ is the resting membrane potential, 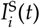 is the total synaptic current onto compartment *i*, and 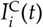 is the current from neighbouring compartments. For neurons with multiple dendritic compartments, we additionally include a voltage-gated calcium-channel current, 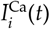, to support active propagation of depolarization towards the soma.

The total synaptic current onto compartment *i*, 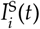, is given by

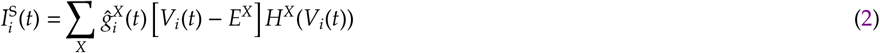

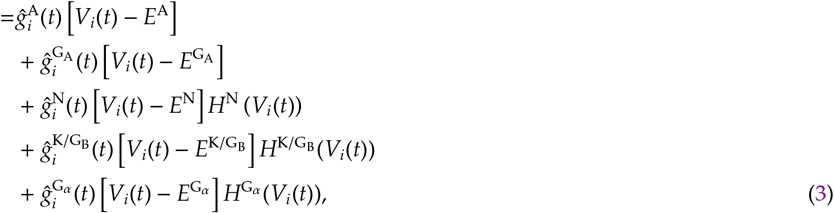

where 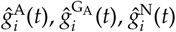, and 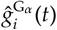 denote the total conductances of AMPA, GABA_A_, NMDA, and *α*5-GABA_A_ channels on compartment *i*, respectively, whereas 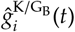 denotes the total conductance of inwardly rectifying potassium (Kir) channels opened through GABA_B_ activation. The corresponding reversal potentials are denoted by 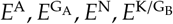, and 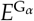. The functions *H*^N^(*V*), 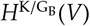, and 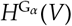 describe the nonlinear voltage dependence of NMDA, Kir/GABA_B_, and *α*5-GABA_A_ currents, respectively, and are given by

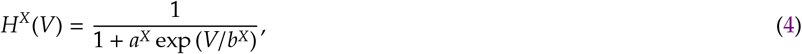

where *a*^*X*^ and *b*^*X*^ are parameters and *X* = {N, K/G_B_, G_*α*_}. For the linear channels, 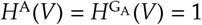 for all *V*.

The temporal evolution of each synaptic conductance is described by

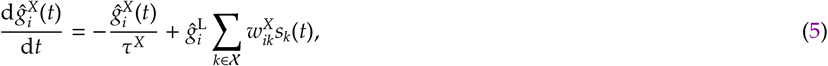

where *τ*^*X*^ is the time constant of channel *X*, 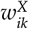 is the connection strength from presynaptic neuron *k* onto compartment *i* for channel *X*, χ is the set of presynaptic neurons that activate channel *X*, and *s*_*k*_(*t*) is the spike train of presynaptic neuron *k*. Here, 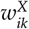 is expressed relative to the leak conductance 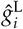. The spike train is defined as

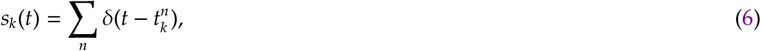

where *δ*(·) is the Dirac delta and 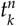 is the time of the *n*th spike of presynaptic neuron *k*. Presynaptic neurons may represent either external input sources or neurons within the simulated recurrent network, as described below.

We define the baseline connection strength from presynaptic neuron *k* to dendritic compartment *i* as 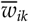. Excitatory connections comprise both AMPA and NMDA components, with 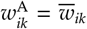 and 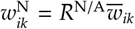, where *R*^N/A^ is the ratio of NMDA to AMPA synaptic strength. We consider two classes of dendrite-targeting inhibitory interneurons in our simulations. A standard inhibitory interneuron, motivated by somatostatin-expressing (SOM) GABAergic cells, comprises both GABA_A_ and Kir/GABA_B_ components, with 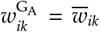 and 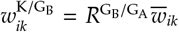, where 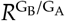 is the ratio of Kir/GABA_B_ to GABA_A_ synaptic strength. A gating-specific interneuron, motivated by nitric-oxide-synthase-expressing (NOS) neurogliaform cells, contributes only an *α*5-GABA_A_ component, with 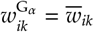.

The current flowing from neighbouring compartments into compartment *i*, 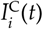, is given by

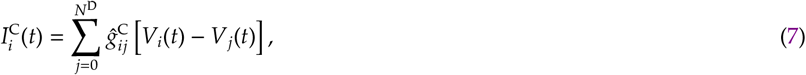

where 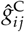 is the passive conductance coupling compartment *j* to compartment *i*. For convenience, we denote the somatic membrane potential by *V*_0_(*t*), so the sum starts at *j* = 0 to include the soma, whose dynamics are defined below.

The current through the VGCC in compartment *i*, 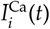, is given by

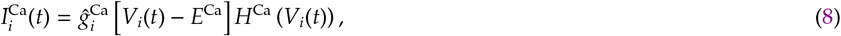

where *E*^Ca^ is the reversal potential of the VGCC, 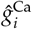 is the corresponding conductance, and *H*^Ca^(*V*) describes its voltage dependence. For simplicity, we use the same functional form for *H*^Ca^(*V*) as for the nonlinear synaptic activation functions (Eq. 4) with a hard activating threshold,

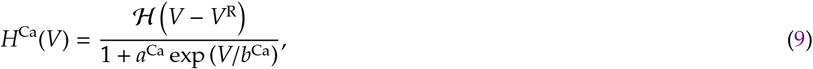

where ℋ(*x*) is the Heaviside step function, with ℋ(*x*) = 1 for *x* ⩾ 0 and ℋ(*x*) = 0 otherwise.

Dividing Eq. 1 by 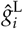 yields the normalized membrane-potential dynamics,

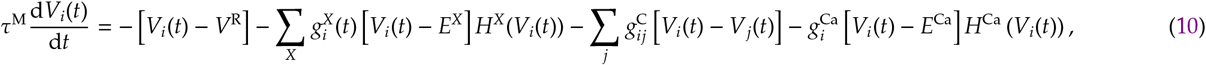

Where 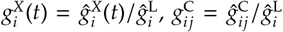, and 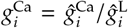 are conductances normalized by the leak conductance and are therefore dimensionless.

#### Somatic compartment

The somatic membrane potential, *V*_0_(*t*), evolves according to

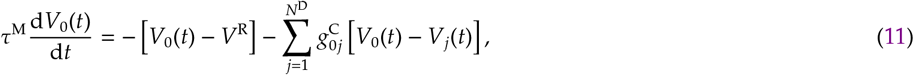

which has the same normalized form as the dendritic dynamics in Eq. 10, but without synaptic or VGCC terms. A neuron emits a spike when the somatic membrane potential, *V*_0_(*t*), crosses the threshold *V*^TH^ from below. After a spike, *V*_0_(*t*) is reset to *V*^RT^ and clamped at that value for the duration of the refractory period, *τ*^R^.

#### Dendritic architecture (morphology)

In our framework, dendritic architecture is specified by the passive coupling matrix 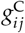, which is zero for uncoupled compartments and equal to *g*^C^ for dendrite-to-soma connections and 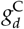 for dendrite-to-dendrite connections. For simplicity, we model passive current flow only from leaf dendrites towards the soma and neglect back-propagation.

#### Single dendrite and two dendrites

For simulations with a single dendritic compartment, we neglect passive current flow from the soma to the dendrite, reflecting the assumption that the total dendritic membrane area is much larger than the somatic area, Δ*A*_dend_ ≫ Δ*A*_soma_. The passive coupling is therefore

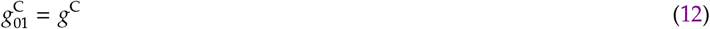

and

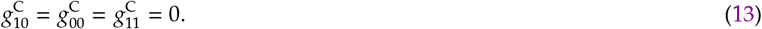

For simulations with two dendritic compartments, we make the same approximation and set

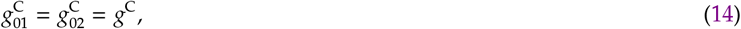

and

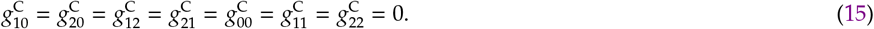

#### Multiple dendrites

For neurons with multiple dendritic compartments, we consider a full binary dendritic tree with *N*^S^ = 2^*L*−1^ leaf dendrites (with *L* representing the number of layers) and a total of *N*^D^ = 2*N*^S^ − 1 dendritic compartments. The first layer contains the *N*^S^ leaf dendrites, the next layer contains *N*^S^/2 dendrites, and so on until the final root compartment. Leaf dendrites are indexed by *i* = 1, …, *N*^S^, the next layer by *i* = *N*^S^ + 1, …, 3*N*^S^/2, and so on. With this indexing convention, passive coupling between dendritic compartments is defined as

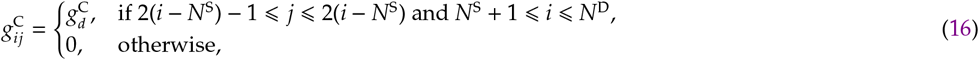

for *i, j* = 1, …, *N*^D^. The soma is then connected to the final dendritic compartment,

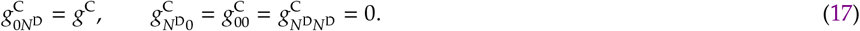

#### External input – single neuron

In single-neuron simulations, we consider two types of external Poisson input: constant firing rate and fluctuating firing rate following a modified Ornstein-Uhlenbeck process. A dendritic compartment receives external synaptic input from presynaptic neurons modelled as point processes, which we refer to as *external neurons*. A total of *N* external neurons connect onto a dendrite, with *N*^E^ excitatory and *N*^I^ inhibitory. The probability that external neuron *k* emits a spike at time *t* is denoted by 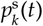. Here, excitatory neurons are defined as 1 ⩽ *k* ⩽ *N*^E^ and inhibitory as *N*^E^ + 1 ⩽ *k* ⩽ *N*. Each spike updates the corresponding synaptic conductance according to Eq. 5. We also impose a refractory period on external neurons in these simulations, such that 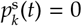 during the refractory interval.

#### Constant firing rate (blanket inhibition)

In this condition, excitatory external neurons always follow the fluctuating Ornstein-Uhlenbeck-based firing-rate process described below. Only inhibitory external neurons fire with a constant rate outside the refractory period. Specifically, for inhibitory neurons,

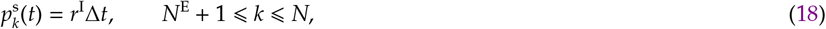

where *r*^I^ is the constant inhibitory firing rate and Δ*t* is the simulation time step.

#### Fluctuating firing rate (matched inhibition)

In this condition, excitatory external neurons follow a fluctuating firing-rate process, and inhibitory external neurons follow the same fluctuations up to a population-specific scaling factor. The spike probability of external neuron *k* is

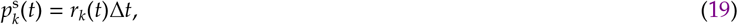

where *r*_*k*_(*t*) is its instantaneous firing rate. We separate the inputs into *N*^OU^ groups with correlated spike patterns. For excitatory inputs, this means *N*^E^*/N*^OU^ neurons per group, and for inhibitory inputs, *N*^I^*/N*^OU^. Each group follows an independent fluctuating firing rate. The instantaneous firing rate is taken as the positive part of an auxiliary variable 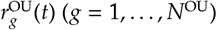 evolving according to

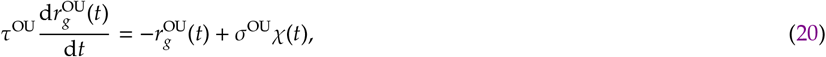

where *τ*^OU^ is the correlation time constant, *σ*^OU^ is the noise amplitude, and *χ*(*t*) is white noise. We then set

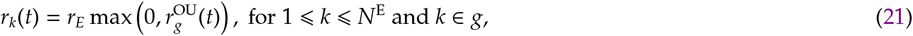

for excitatory inputs, and

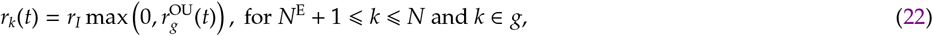

for inhibitory inputs. As in the blanket-inhibition condition, all external neurons obey a refractory period.

#### External input – recurrent network

In recurrent-network simulations, external input is used either to activate a memory from a silent initial condition, to switch recall between memories, or to gate dendrites through targeted inhibitory input. In all cases, spikes follow a homogeneous Poisson process without a refractory period, reflecting the approximation that many external neurons contribute simultaneously and thereby reducing computational cost. The probability that an external spike arrives onto dendrite *k* of neuron *i* is

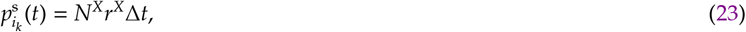

where *N*^*X*^ is the assumed number of external neurons of type *X* and *r*^*X*^ is their average firing rate.

#### Short burst for memory activation

To activate a memory in the recurrent network, we apply a brief burst of external excitatory input after an initialisation period. In single-dendrite networks, no dendritic index is required. In multiple-dendrite networks, the externally activated dendrite is specified explicitly. The set of externally activated neurons, and dendrites when applicable, is denoted by ℰ. For all *i*_*k*_ ∈ ℰ, the probability that dendrite *k* of neuron *i* receives an external spike is given by 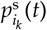, as defined above.

#### Constant firing rate for inhibitory gating

In simulations with two-dendrite neurons and external inhibitory gating, each neuron has two dendrites connected to the soma. To gate one memory set, we select one dendrite index, *k* ∈ {1, 2}, across the entire network. For the selected dendrite of each neuron, external inhibitory spikes drive the *α*5-GABA_A_ conductance with probability 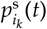, as defined above.

#### Connectivity – single neuron

In the single-neuron simulations, the ratio of excitatory to inhibitory external neurons is fixed at 4 : 1. All external neurons connect to the postsynaptic neuron with uniform synaptic weights, 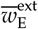 for excitatory input and 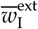 for inhibitory input.

#### Connectivity – recurrent network

In all recurrent-network simulations, synaptic connectivity is first generated in the binary model using the modified perceptron learning rule and then mapped onto the biophysical model, as described below. The different recurrent-network configurations considered here differ only in how the mapped connectivity is distributed across dendrites and in how external stimulation or gating is applied.

#### Single memory matrix

For networks with one dendrite per neuron, the mapped connectivity matrix is assigned to that dendrite. A subset of neurons receives external excitatory input during a brief stimulation period, with synaptic weight 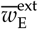.

#### Two memory sets with optional external gating

For networks with two dendrites per neuron and two memory sets, we generate two connectivity matrices from two independently generated sets of memory patterns and assign one matrix to each dendrite. External excitatory input is then applied during a brief stimulation period to a selected set of neurons, targeting one or both dendrites, with synaptic weight 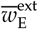. When external inhibitory gating is included, the selected dendrite on each neuron additionally receives inhibitory input during the gating period, with synaptic weight 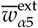^t^.

#### Autonomous readout-gating architecture

For the autonomous configuration, we simulate three interconnected networks: a memory network, a readout network, and a gating network. In the memory network, neurons have multiple dendrites, and each dendrite is assigned a connectivity matrix encoding a different memory set. In the readout network, neurons have a single dendrite; excitatory readout neurons receive input from excitatory neurons in the memory network with connections generated using the modified perceptron learning rule. Recurrent readout connectivity is generated using the modified perceptron learning rule as well. The gating network consists of inhibitory neurons that receive input from excitatory readout neurons and project back to the dendrites of the memory network.

#### Simplified fluorescence signals (ΔF/F)

To relate our results to standard experimental measurements, we implement a simplified normalized fluorescence signal as a slow and noisy proxy for large dendritic depolarizations in calcium imaging^32^. For simplicity, we use the NMDA voltage-dependent nonlinearity, *H*^N^(*V*_*i*_(*t*)), as the nonlinear activation term driving the fluorescence signal. We model these dynamics as a nonlinear voltage-dependent signal followed by low-pass filtering and additive noise,

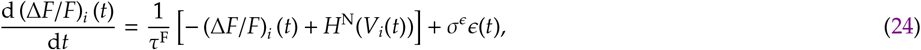

where (Δ*F/F*)_*i*_(*t*) is the normalized fluorescence of compartment *i* at time *t, τ*^F^ is the effective fluorescence time constant and *ϵ*(*t*) is a white noise term with amplitude *σ*^*ϵ*^.

### Analyses of the biophysical model

#### Dendritic compartment fixed points

To analyse the fixed points of the dendritic membrane-potential dynamics, we approximate the synaptic conductances using average presynaptic firing rates. For simplicity, we suppress the dendritic-compartment index whenever no ambiguity arises, for example 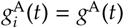 and 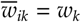. The temporally averaged conductance of channel *X* is then

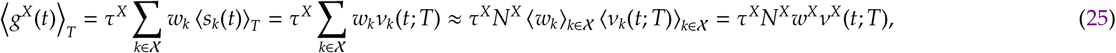

where the temporal average over a window of duration *T* is defined as

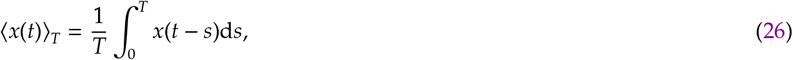

and *ν*_*k*_(*t*; *T*) = ⟨*s*_*k*_(*t*) ⟩_*T*_ denotes the temporally averaged firing rate of presynaptic neuron *k*. Here, *N*^*X*^ is the number of presynaptic neurons whose transmitter activates channel *X*. The corresponding population averages are

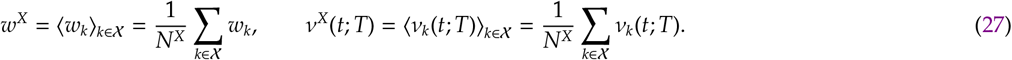

Taking the temporal average of the membrane-potential dynamics (Eq. 10) and considering an isolated dendritic compartment, we approximate the nonlinear current terms by evaluating them at the temporally averaged membrane potential. This yields

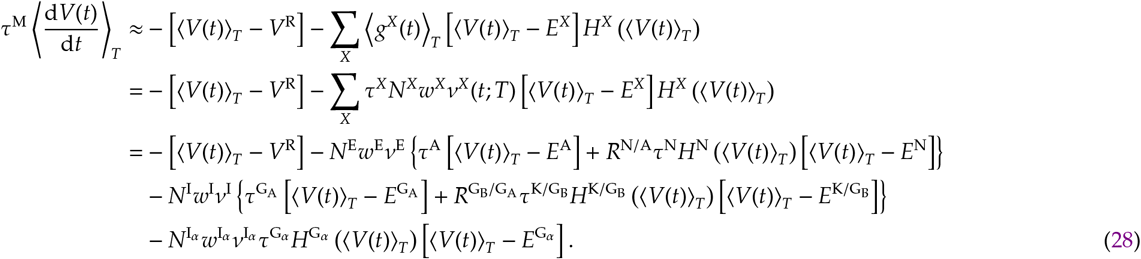

Setting Eq. 28 to zero allows us to evaluate numerically the fixed-point voltages, *V*^∗^, as a function of excitatory and inhibitory input, thereby defining a state space. Without *α*5-GABA_A_ gating inhibition—i.e., with 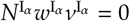 in Eq. 28—we obtain

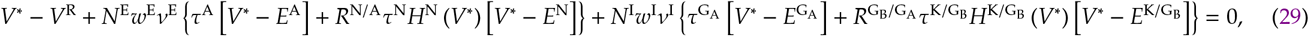

which can be rewritten as

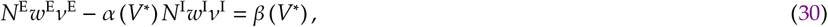

where

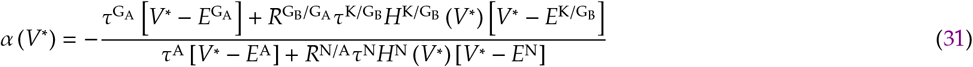

and

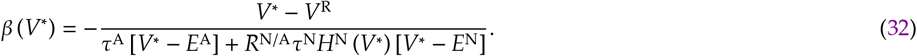

Eq. 30 defines the relationship between excitatory input, *N*^E^*w*^E^*ν*^E^, and inhibitory input, *N*^I^*w*^I^*ν*^I^, required for a fixed point at voltage *V*^∗^. In this expression, *α*(*V*^∗^) is dimensionless and sets the relative weighting of inhibition with respect to excitation, whereas *β*(*V*^∗^) has units of frequency, like *ν*^E^ and *ν*^I^, because synaptic weights are dimensionless. The resulting phase diagram contains three regions: one with a single stable hyperpolarised fixed point, one with a single stable depolarised fixed point, and one in which stable hyperpolarised and depolarised fixed points coexist with an unstable fixed point.

For the associative memory network, we are particularly interested in the line corresponding to the fixed point *V*^∗^ = *V*^min^, where *V*^min^ is the least depolarised dendritic membrane potential that still elicits somatic spiking. This line is described by

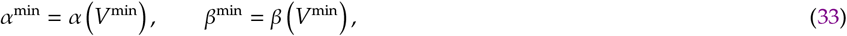

where *V*^min^ is calculated below.

#### Postsynaptic neuron’s firing rate

We first calculate the least depolarised dendritic membrane potential required to elicit a somatic spike, denoted by *V*^min^. For a soma coupled to a single dendrite with passive coupling *g*^C^, this corresponds to the minimum dendritic membrane potential for which the somatic membrane potential reaches the spike threshold at steady state, yielding

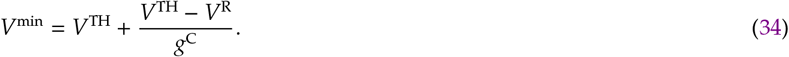

To estimate the postsynaptic firing rate, we calculate the inter-spike interval, *t*^ISI^, assuming that the soma receives constant input from a single dendrite held at the depolarised fixed point, *V*^∗^. Dropping the somatic index for simplicity, such that *V*_0_(*t*) = *V*(*t*), the postsynaptic firing rate is

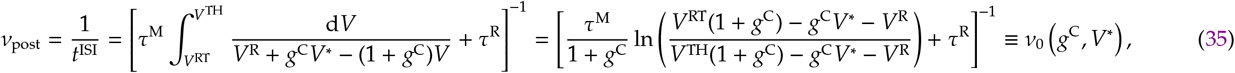

where *ν*_0_(*g*^C^, *V*^∗^) denotes the firing rate of a neuron with a single dendrite connected to the soma by passive coupling *g*^C^ and held at a constant dendritic membrane potential *V*^∗^.

#### Minimum GABA_A_ conductance

Excitatory and inhibitory currents balance most effectively when GABA_A_ conductance remains continuously above zero. To quantify this, we define a practical lower-bound estimate of the GABA_A_ conductance as the temporal mean minus half a standard deviation, evaluated for Poisson input. For a Poisson process, the variance of the conductance is

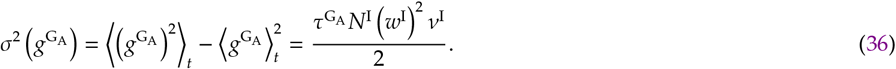

Using the temporal mean conductance from Eq. 25, we then define

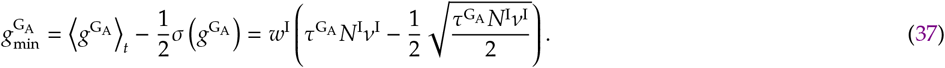

#### Interspike interval distribution

We estimate the distribution of network interspike intervals (ISIs) from the firing rate of active neurons and the timing of memory transitions. For externally driven recall, memories are activated periodically with period *T*. Assuming that neurons active during recall fire at rate *ν*_0_ (Eq. 35), the ISI distribution, *t*_ISI_, is

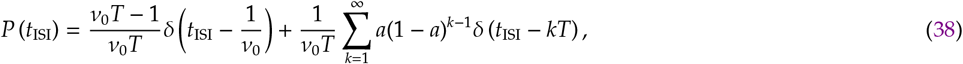

where *a* is the activity level of a memory pattern. The first term corresponds to successive spikes emitted while the neuron remains active within a recalled memory, whereas the second term accounts for intervals spanning one or more memory periods between successive recall events. In Fig. 2G, the theoretical prediction is converted to spike counts using the same binning as in the simulations. The mean of the distribution is

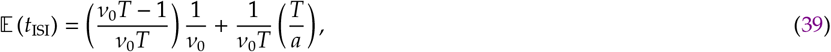

and the second moment is

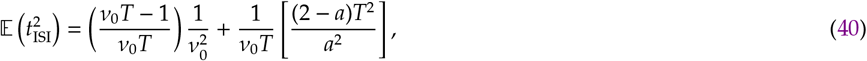

which together yield the coefficient of variation,

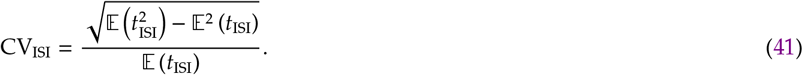

### Binary model

#### Network dynamics

For clarity, we refer to binary neurons as *units*, which take values of zero or one. The state of unit *i* evolves in discrete time according to

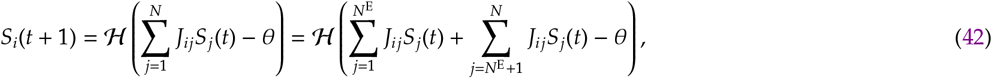

where *S*_*i*_(*t*) is the state of unit *i* at discrete time *t*, and *J*_*ij*_ is the connection strength from presynaptic unit *j* to postsynaptic unit *i*. The variables *N*^E^ and *N*^I^ denote the numbers of excitatory and inhibitory units, respectively, such that *N* = *N*^E^ + *N*^I^ is the total number of units. The threshold *θ* is positive (*θ* ⩾ 0), and ℋ (*x*) is the Heaviside step function, with ℋ(*x*) = 1 for *x* ⩾ 0 and ℋ(*x*) = 0 otherwise (note the repeated definition for clarity). In Eq. 42, the input is separated into excitatory and inhibitory components according to the sign of the outgoing connections,

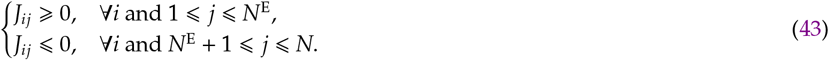

#### Memory patterns

We denote memory patterns by 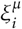, where 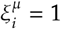 indicates that unit *i* is active in memory *µ*, and 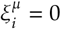 indicates that it is inactive. Thus, 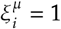 means that unit *i* belongs to the assembly, or engram, representing memory *µ*. We consider both independent and correlated sets of memory patterns.

#### Independent memory patterns

Independent memory patterns are generated as

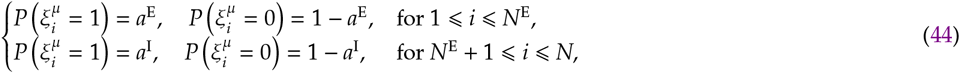

where *a*^E^ and *a*^I^ are the probabilities that excitatory and inhibitory units, respectively, are active in a given memory pattern.

#### Correlated memory sets

We also consider correlated memory sets. To do so, we generate one set of independent memory patterns, as described above, and additional sets whose patterns are correlated with those in the reference set. For two patterns 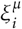 and 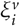 with *µ* ≠ *ν*, their correlation is

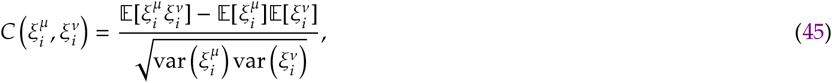

where 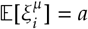 is the activity level and 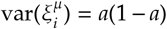 is the corresponding variance. For notational simplicity, we omit the excitatory and inhibitory superscripts here, although the implementation still uses distinct activity levels *a*^E^ and *a*^I^. Because the activity level is fixed, 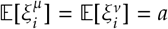 and 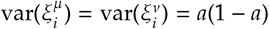. Moreover, 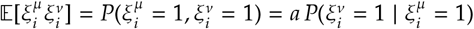. Substituting these expressions into Eq. 45 gives

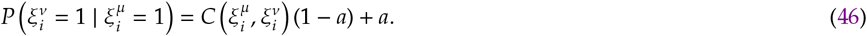

Using the symmetry 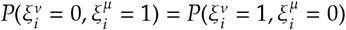 together with the complement relation 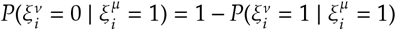 yields

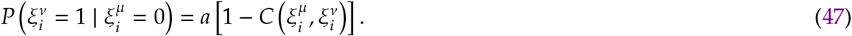

We use these conditional probabilities to generate correlated patterns. Starting from a random pattern 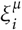 sampled according to Eq. 44, we set 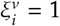 with probability 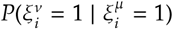 if 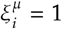, and with probability 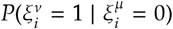 if 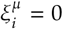; otherwise, we set 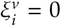. In the implementation, Eqs. 46 and 47 are applied separately to excitatory and inhibitory units using *a*^E^ and *a*^I^, respectively.

#### Readout

For simplicity, we denote the memory patterns of the readout network by 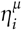. The readout network represents all memories stored in the memory network, indexed by *µ* = 1, …, *N*^S^*p*, where *N*^S^ is the number of memory sets (and dendritic leaves) in the memory network and *p* is the number of memory patterns per set. Excitatory readout units encode these memories without overlap,

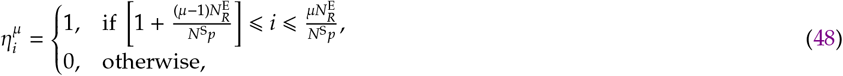

where 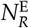 is the number of excitatory readout units. Inhibitory readout units encode memories randomly according to

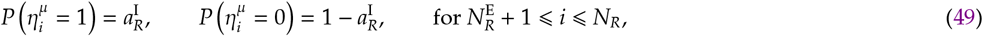

where 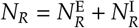 is the total number of readout units and 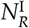 is the number of inhibitory readout units.

#### Gating

For simplicity, we denote the activity patterns of the gating network by 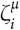. The gating network contains *N*_*G*_ inhibitory units that target dendrites in the memory network. We assume a binary dendritic tree, so that the number of memory sets (equivalently, leaf dendrites) satisfies *N*^S^ = 2^*L*^ for some integer *L*. Under this architecture, each neuron has 2*N*^S^ − 2 dendrites that require inhibitory gating (excluding the root compartment directly connected to the soma) corresponding to the number of gating patterns, *µ* = 1, …, 2*N*^S^ − 2.

These patterns are non-overlapping and are defined as

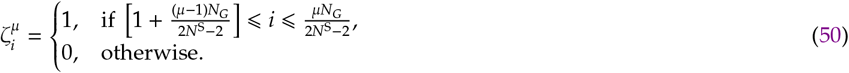

#### Connectivity and connection weights

Connections in the recurrent-network simulations are generated from memory patterns in two ways. Some weights are optimized using a modified Perceptron Learning Rule (modPLR), whereas others are assigned directly from the structure and overlap of the corresponding coding patterns. We describe these two constructions below.

#### Recurrent memory-encoding weights

We choose the network weights of excitatory and inhibitory units such that the memory patterns 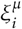 of a memory set, *µ* = 1, …, *p*, are fixed points of the binary dynamics (Eq. 42). To quantify this, we use the stability parameter^97^, 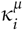, which for unit *i* and memory *µ* is defined as

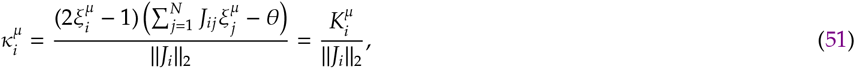

where 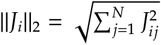 is the *ℓ*_2_ norm of the incoming weight vector to unit *i*. If 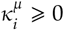 for all *i* in a given memory pattern 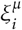, then that pattern is a fixed point of the network dynamics, as follows directly from Eq. 42.

For all patterns in a memory set to be fixed points of the dynamics, we focus first on the numerator of Eq. 51, because the denominator affects the magnitude of the stability parameter but not whether a pattern is itself a fixed point. To do so, we use a modified Perceptron Learning Rule (modPLR), adapted from Brunel ^17^ . The modPLR reformulates the numerator of Eq. 51 into two state-dependent constraints,

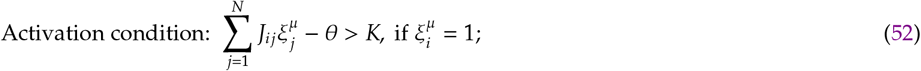

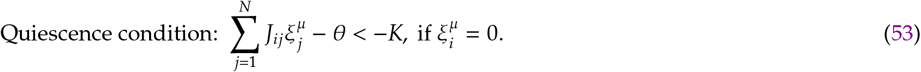

For a given memory pattern *µ* and unit *i*, the modPLR then applies the following update rules:

**Step 1**. Increase the unmasked input connections (for which mask_*ij*_ = 0) by *λ* if the activation condition (Eq. 52) is violated.

**Step 2**. Decrease the unmasked input connections (for which mask_*ij*_ = 0) by *λ* if the quiescence condition (Eq. 53) is violated.

**Step 3**. Enforce Dale’s law for excitatory units: if 1 ⩽ *j* ⩽ *N*^E^ and *J*_*ij*_ *<* 0, set *J*_*ij*_ = 0.

**Step 4**. Enforce Dale’s law for inhibitory units: if *N*^E^ *< j* ⩽ *N* and *J*_*ij*_ *>* 0, set *J*_*ij*_ = 0.

Initially, we set mask_*ij*_ = 0 for all *i, j*. A single optimisation iteration consists of repeated passes through all units and memory patterns until the update conditions above are satisfied. After each such iteration, connections that have been eliminated are permanently masked by setting mask_*ij*_ = 1 whenever *J*_*ij*_ = 0. Importantly, a connection may temporarily reach zero within an iteration as long as it remains unmasked, i.e., as long as mask_*ij*_ = 0.

Because the stability parameter in Eq. 51 is normalised by ∥*J*_*i*_∥_2_, stability can be improved by reducing the *ℓ*_2_ norm of the incoming weight vector. However, biological connectivity, for example in hippocampal CA3, is also sparse^22^, which motivates reducing the *ℓ*_1_ norm, 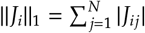, as well. To promote both effects, after convergence of the modPLR update rules we apply the shrinkage step

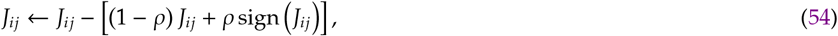

where *ρ* ∈ [0, 1] controls the relative contributions of the *ℓ*_2_- and *ℓ*_1_-norm penalties (see Supplementary Text for the derivation). This shrinkage step generally renders some memories unstable and therefore initiates a new optimisation iteration. The procedure is repeated until 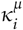 converges (Fig. S3). The optimisation is initialized with weights drawn independently from a uniform distribution, *J*_*ij*_ ∼ 𝒰 (−*J*^∗^, *J*^∗^), where *J*^∗^ is a free parameter.

#### Readout weights

Readout-network weights are generated using the same modified perceptron learning rule (modPLR) used for the memory network, with the addition that excitatory readout units also receive input from the memory network. Accordingly, the activation and quiescence conditions are rewritten as

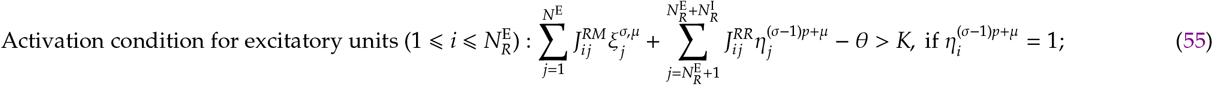

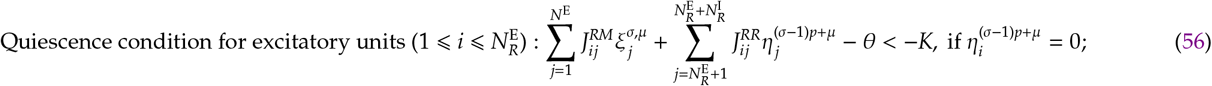

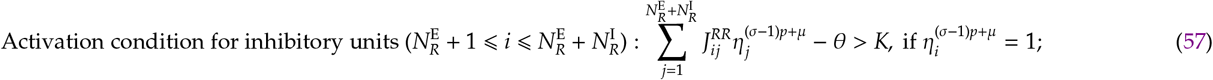

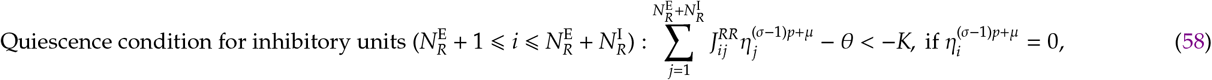

where 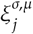 denotes the activity of memory-network unit *j* in memory *µ* of memory set *σ*, 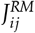 is the connection from memory-network unit *j* to readout-network unit *i*, and 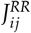 is a recurrent connection within the readout network. We then optimize both 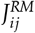 and 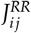 using the same modPLR procedure described above for the memory network.

#### Readout-to-gating network weights

Because excitatory readout units encode individual memories and inhibitory gating units encode dendritic branches, we connect these two populations so that readout activity associated with memory set *σ* activates the gating units corresponding to the same set. We define these weights as

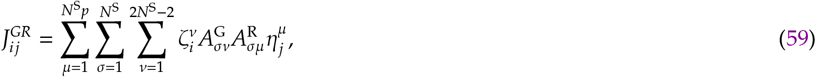

where 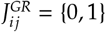 is the connection from presynaptic readout unit *j* to postsynaptic gating unit *i*. To specify the correct readout-to-gating connectivity, we introduce two auxiliary matrices, 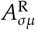 and 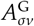. Here, 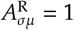 if readout memory *µ* belongs to memory set *σ*, and 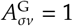 if gating pattern, and thus dendrite, *ν* belongs to memory set *σ*. Because the readout memories are ordered as non-overlapping blocks (Eq. 48), the auxiliary matrix 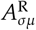 is given by

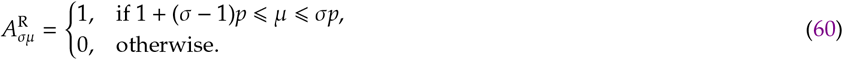

The auxiliary matrix 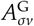 depends on the dendritic-tree architecture and indexing convention defined above (Eq. 16). In the memory network, each neuron has 2*N*^S^ − 2 gated dendrites, indexed by *ν* = 1, …, 2*N*^S^ − 2, where the *N*^S^ leaf dendrites occupy the first layer and each subsequent layer contains half as many dendrites. The matrix 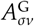 specifies whether dendrite *ν* belongs to the path associated with memory set *σ*. It is given by

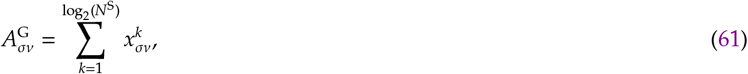

where

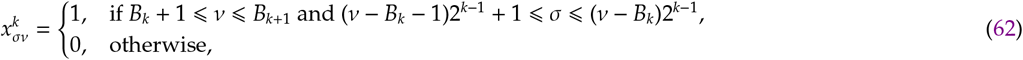

with

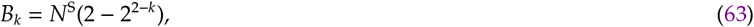

which denotes the total number of dendrites in all layers below layer *k*. The sum in Eq. 61 runs over dendritic layers and evaluates to one only when the conditions in Eq. 62 are satisfied. The first condition identifies the layer containing dendrite *ν*, and the second identifies the memory sets *σ* associated with that dendrite.

#### Gating weights

Following the dendritic-tree architecture and indexing defined above, we denote *d* = 1, …, 2*N*^S^ −2 as the index of a gated dendrite across all layers. The weight matrix connecting gating unit *j* to dendrite *d* is then defined as

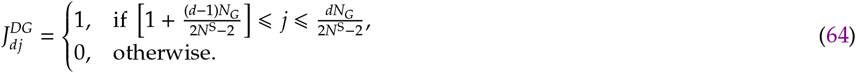

### Binary to biophysical mapping

To map the binary network onto the biophysical network, we match the thresholded input of a binary unit to the thresholded input of a biophysical neuron evaluated at the minimum active fixed point (Eqs. 30 and 33). We assume that both excitatory and inhibitory neurons fire at a common rate *ν*^∗^ when their dendrites are in the active depolarised state. Under this approximation, the firing-rate dynamics of the biophysical network can be written as

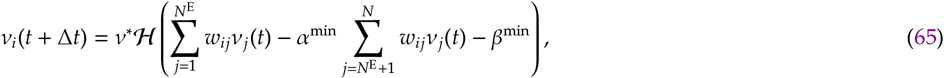

where *α*^min^ and *β*^min^ are defined in Eqs. 30–33, and *ν*^∗^ is the postsynaptic firing rate given by Eq. 35. Here, *w*_*ij*_ denotes a non-negative biophysical synaptic strength, with inhibitory contributions represented by the explicit prefactor *α*^min^ in Eq. 65.

For a memory pattern *µ*, and dropping the time index as well as the superscript “min” for readability, Eq. 65 becomes

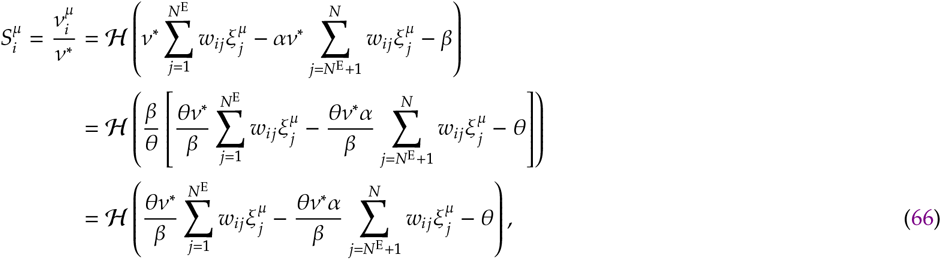

where 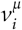 is the firing rate of neuron *i* when all neurons belonging to memory pattern *µ* fire at rate *ν*^∗^ and all other neurons are silent, so that 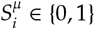 as in the binary model. Because *θ >* 0 and *β >* 0 in the regime considered here, the identity ℋ (*gx*) = ℋ (*x*) for *g >* 0 is valid in the second equality of Eq. 66.

Matching the arguments of the Heaviside functions in Eqs. 42 and 66 yields

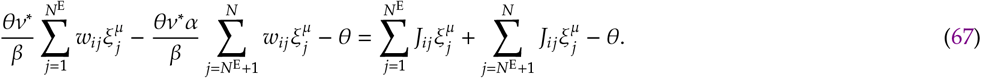

This gives the mapping from binary to biophysical connection weights,

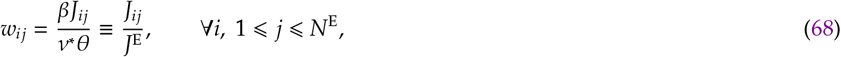

and

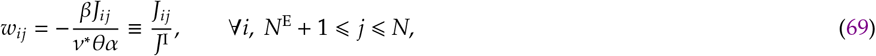

where

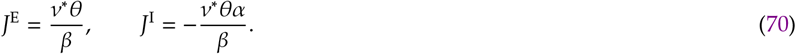

For the memory-to-readout connections, we used the same mapping as above. For the readout-to-readout connections we used the same mapping as above, but scaling the weights to protect memory sets from interference,

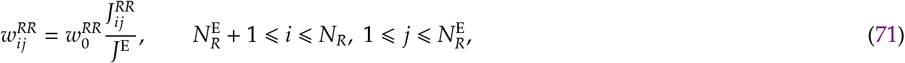

and

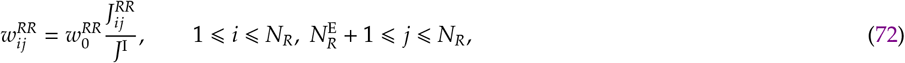

where 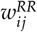 is the weight between neurons *j* and *i* in the readout network, and 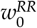 is a scaling parameter. For the readout-to-gating connections, we used

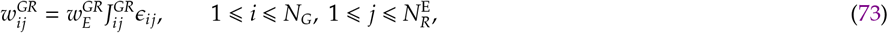

and

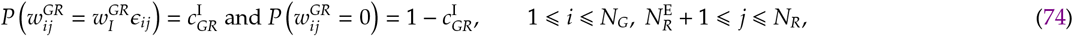

where 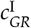 is the probability of a connection from a readout inhibitory neuron to a gating neuron, 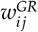 is the weight between neurons *j* from the readout network and *i* from the gating network, 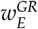 and 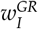 are weight mappings, and *ϵ*_*ij*_ is a random factor drawn uniformly between *ϵ*^∗^ and *ϵ*^∗^ + 1. Note that excitatory and inhibitory connections use different noise levels, 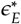 and 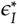, respectively. For gating-to-memory connections, we used

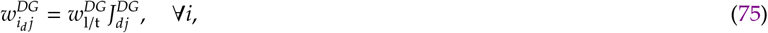

where 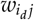 is the connection from gating neuron *j* to dendrite *d* of memory neuron *i* and 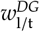 is the weight mapping (l corresponds to a leaf branch and t to a trunk branch).

### Quantifications

#### Correlation measures

We quantify the temporal correlation, *C*(*x, y*), between two variables *x* and *y* using the Pearson correlation coefficient,

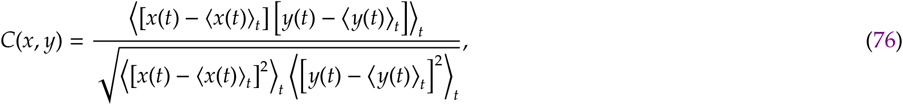

where ⟨·⟩_*t*_ denotes the temporal average.

#### Excitatory-inhibitory correlation

For the excitatory-inhibitory correlation, we compute dendrite-specific correlations using Eq. 76 with

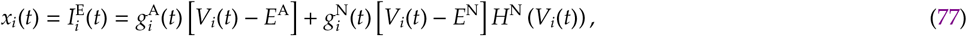

and

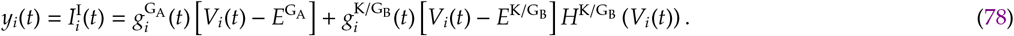

Because excitatory and inhibitory currents have opposite signs, EI balance is reflected by a negative correlation between 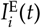 and 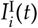.

#### Voltage and ΔF/F correlations

For voltage and fluorescence correlations, we compare different dendrites of the same neuron. In these comparisons, *x*(*t*) denotes the ungated leaf dendrite through which memories are accessible (the recall leaf dendrite), whereas *y*(*t*) denotes either the gated leaf dendrite or the isolated leaf dendrite, which is ungated but lacks somatic access.

#### Memory overlap

To assess whether a given memory is being recalled, we compute its overlap with the network activity using the Pearson correlation coefficient. This metric indicates whether the network activity is correlated (*m*^*µ*^ ≈ 1), anti-correlated (*m*^*µ*^ ≈ −1), or uncorrelated (*m*^*µ*^ ≈ 0) with a target memory pattern *µ*.

#### Time-varying overlap

For simulations involving multiple stimulated memories, the overlap between network activity and memory pattern *µ* at time *t*, denoted by *m*^*µ*^(*t*), is defined as

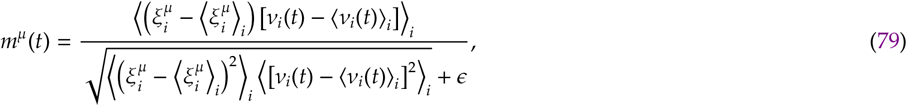

where *ϵ* = 10^−8^ is a small constant used to ensure that the overlap remains zero for inactive networks. Here, ⟨·⟩_*i*_ denotes the average over neurons in the population and *ν*_*i*_(*t*) denotes the number of spikes emitted by neuron *i* within a temporal window of duration *T*^bin^ centred at time *t*,

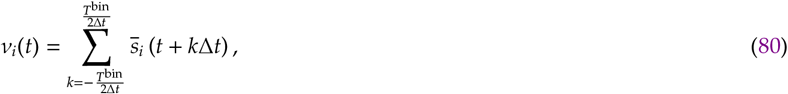

where 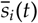 is one if neuron *i* spiked in discrete time *t* (and zero otherwise) given a time-step Δ*t*. A value of *m*^*µ*^(*t*) = 1 indicates that the network activity is perfectly correlated with memory pattern *µ*, whereas *m*^*µ*^(*t*) = 0 indicates that they are uncorrelated at time *t*.

#### Fixed-point overlap

For simulations estimating the basin of attraction, the overlap is computed after a transient period *t*^∗^. We use the same Pearson-based definition as above, but omit the explicit time dependence,

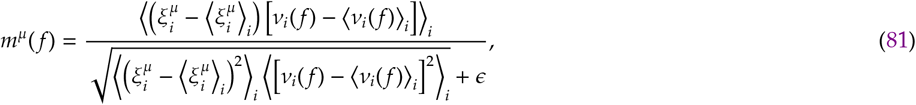

where *ν*_*i*_( *f* ) is the number of spikes emitted by neuron *i* in the interval (*t*^∗^, *t*^∗^ + *T*^m^) when a fraction *f* of memory neurons is externally activated at the start of recall. The time *t*^∗^ accounts for transient dynamics, whereas *T*^m^ defines the integration window used to compute the average fixed-point activity.

#### Reliability

We summarize recall performance, or equivalently the basin of attraction, by the reliability

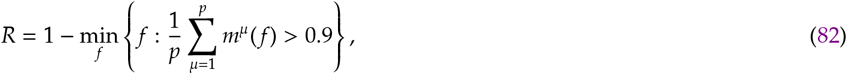

where *m*^*µ*^( *f* ) is the fixed-point overlap (Eq. 81) and *f* is the fraction of externally activated memory neurons, which we refer to as the initial condition size. Reliability is thus one minus the smallest initial condition size for which the average overlap exceeds 0.9: *R* = 1 indicates that recall succeeds without any external activation, whereas *R* = 0 indicates that all memory neurons must be activated.

### Parameters

All parameter values used in the simulations and analyses are listed in Tables S1, S2, S3, and S4. For single dendrite simulations (Figs. 1E– I and S1) we did a parameter sweep over excitatory and inhibitory weights, as well as inhibitory firing-rates to find combinations of parameters for which the postsynaptic neuron fired at approximately a target single-neuron firing rate, denoted by *ν*^single^.

## Supplementary figures

**FIG. S1.**
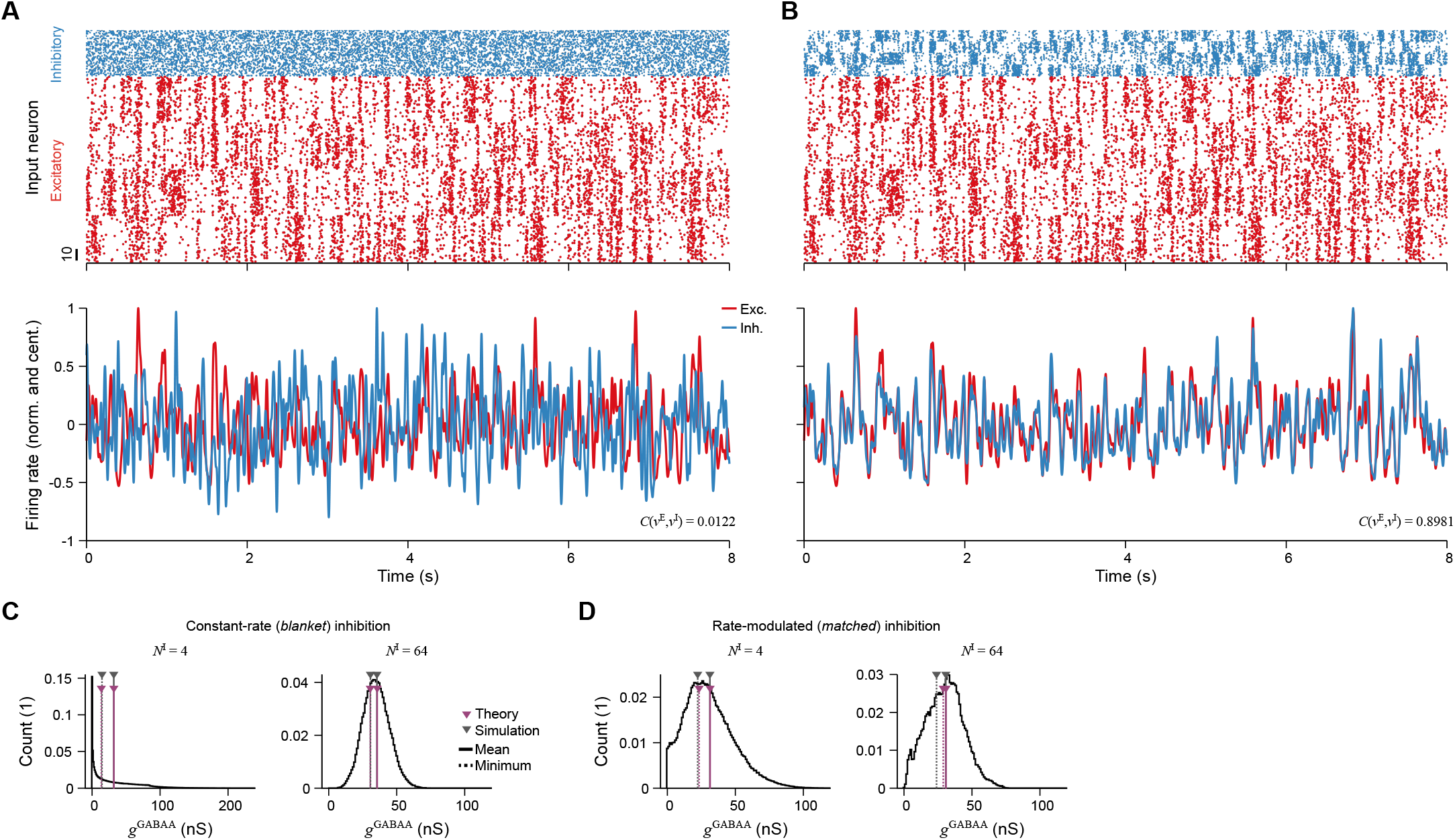
Input statistics and inhibitory conductance for the single-neuron simulations. **A**, Raster plot of excitatory and inhibitory spikes received by the dendrite of a single neuron (top) and the corresponding normalised firing rates centred at zero (bottom). Excitatory spikes follow a modified Ornstein-Uhlenbeck process that is independent across four neuron groups (see Methods). Inhibitory spikes fire at a constant rate, corresponding to blanket inhibition (see Methods). The correlation between excitatory and inhibitory firing rates is denoted by *C*(*ν*^E^, *ν*^I^). **B**, Same as panel A, but inhibitory spikes follow the same spike probabilities as the excitatory spikes, corresponding to matched inhibition (see Methods). **C**, Distribution of GABA_A_ conductance for simulations with *N*^I^ = 4 (left) and *N*^I^ = 64 (right) inhibitory inputs under blanket inhibition (cf. Fig. 1Ei, bottom). Vertical lines with arrowheads indicate the mean conductance and the estimated minimum GABA_A_ conductance from theory and simulation. **D**, Same as panel C, but for matched inhibition (cf. Fig. 1Eii, bottom).

**FIG. S2.**
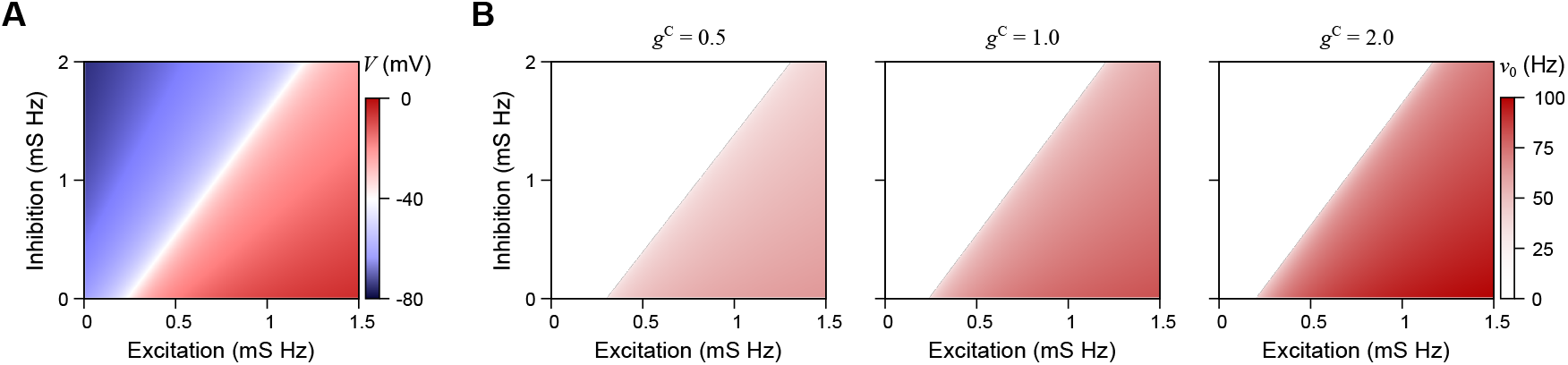
Postsynaptic firing rate derived from membrane-potential fixed points. **A**, Membrane-potential fixed points (colour) as a function of excitatory and inhibitory input (same as Fig. 1D, left). **B**, Postsynaptic firing rate (colour) as a function of excitatory and inhibitory input for three values of *g*^C^. Firing rates are computed from the membrane-potential fixed points shown in panel A.

**FIG. S3.**
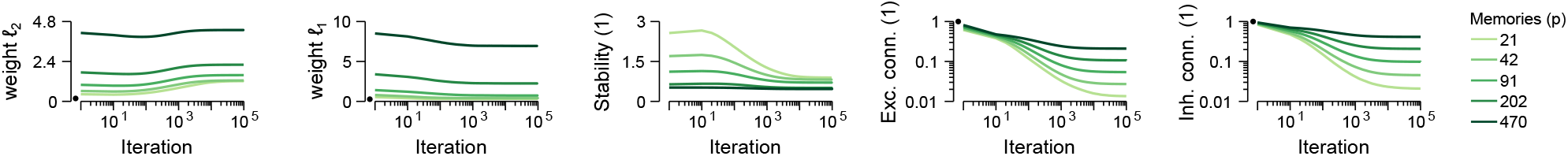
Evolution of binary-network connectivity, weight norms, and stability under the modified perceptron learning rule. Binary-network quantities as a function of iteration number under the modified perceptron learning rule. From left to right: weight *ℓ*_2_ norm, weight *ℓ*_1_ norm, memory-pattern stability, excitatory connectivity, and inhibitory connectivity. Each line corresponds to a different number of stored memories (see legend). Circles indicate initial condition values.

**FIG. S4.**
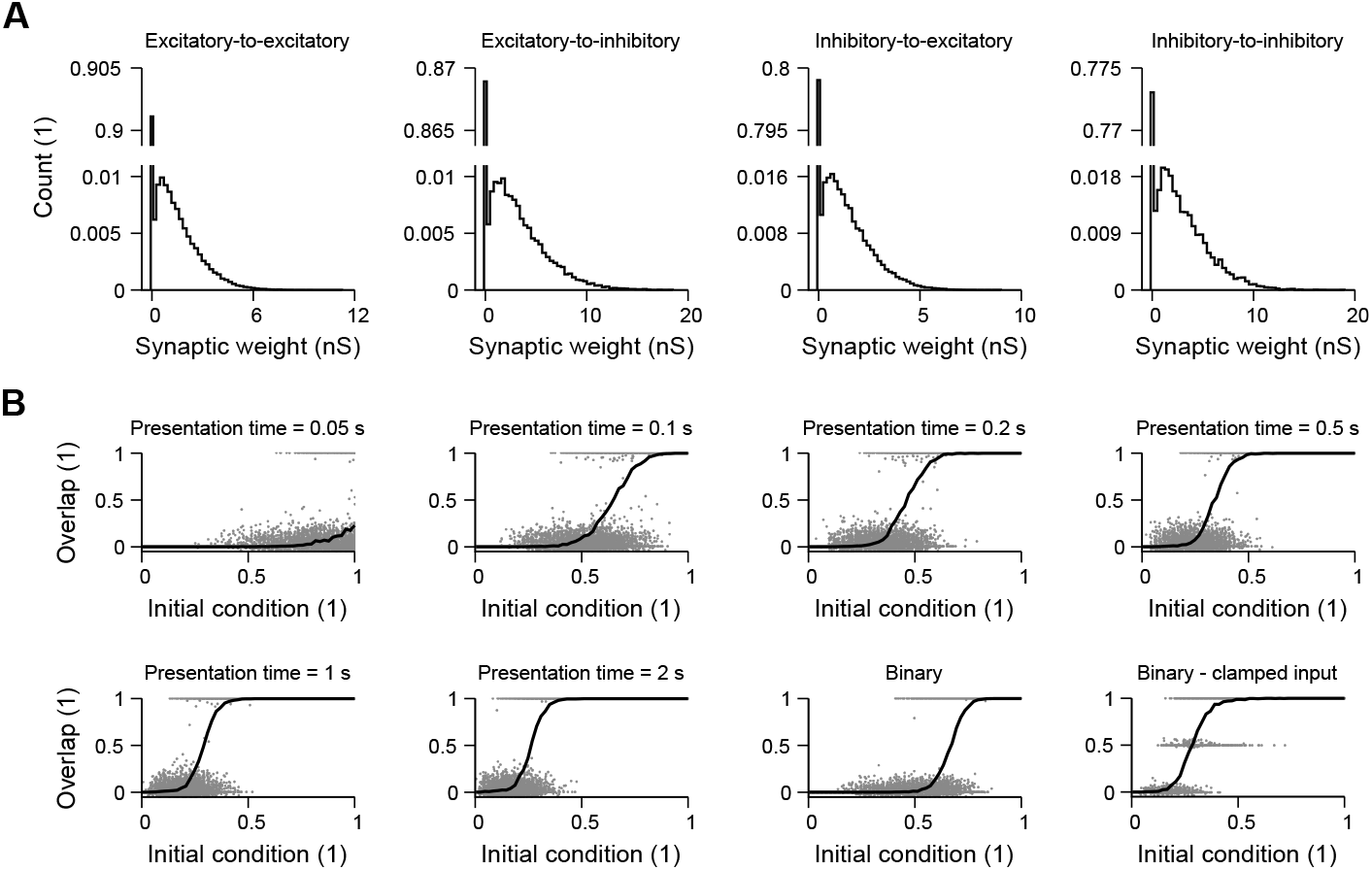
Weight distributions and recall performance in the memory network of Fig. 2. **A**, Distributions of excitatory-to-excitatory, excitatory-to-inhibitory, inhibitory-to-excitatory, and inhibitory-to-inhibitory synaptic weights. Weights are shown in nS, corresponding to the biophysical implementation *w*_*ij*_ (Eq. 5) multiplied by the leak conductance. **B**, Overlap between the final network state and the target memory pattern as a function of initial condition size, defined as the fraction of the memory pattern that is externally activated. Individual points correspond to different memory patterns and distinct groups of activated neurons. The solid line denotes the average (also shown in Fig. 2J).

**FIG. S5.**
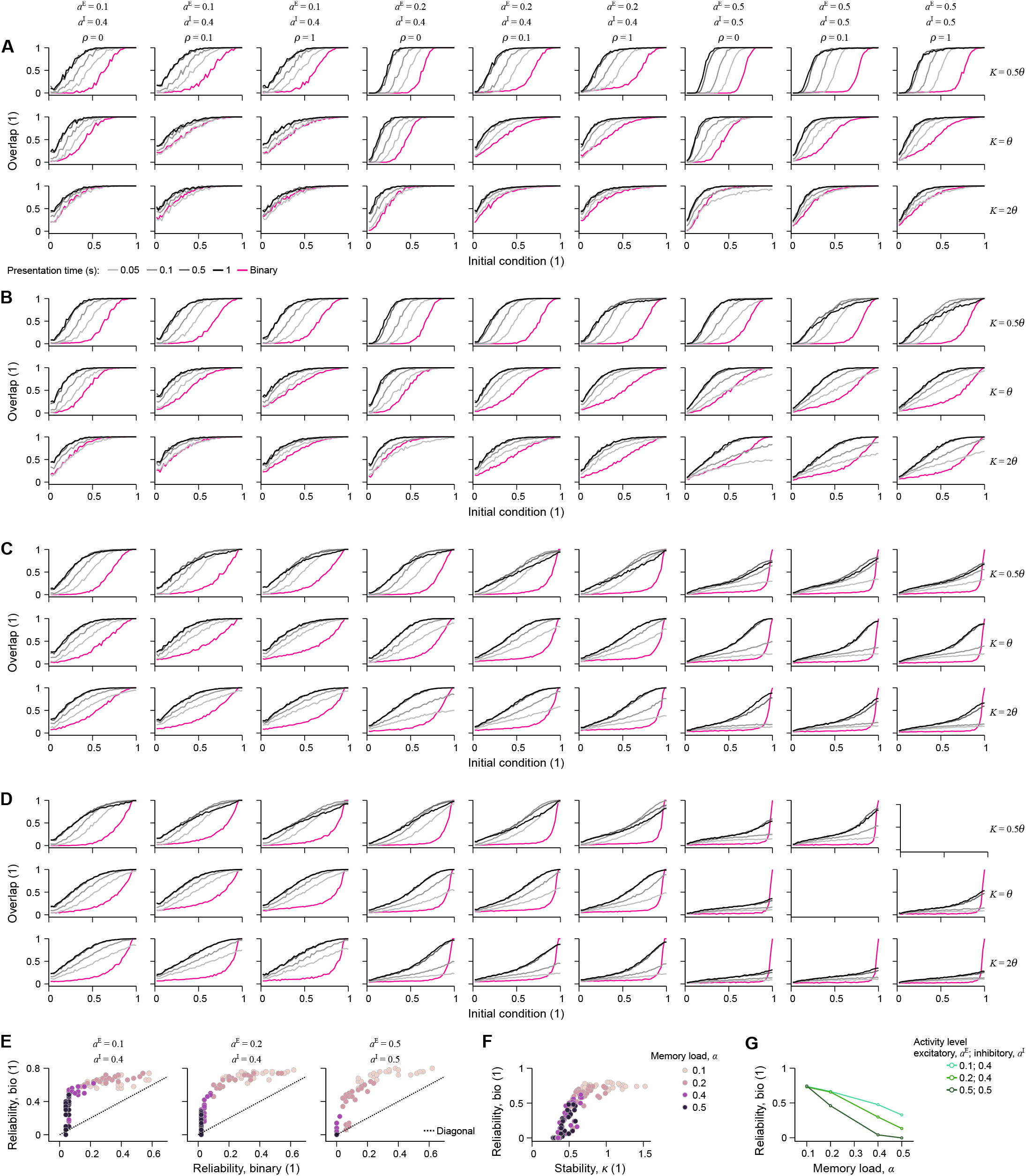
Robustness of biophysical recall across network parameters. Memory-pattern reconstruction, quantified by reliability, is shown for different activity levels *a*^E^ and *a*^I^, regularisation strengths *ρ* (Eq. 54), robustness thresholds *K*, and numbers of stored memory patterns *p*, for networks of size *N* = 100 with *N*^E^ = 80 and *N*^I^ = 20. **A**, Overlap (cf. Fig. 2J) as a function of initial condition size, defined as the fraction of externally activated memory neurons, for networks trained to store *p* = 10 memories (*α* = *p/N* = 0.1). **B**, Same as panel A for networks trained to store *p* = 20 memories (*α* = 0.2). **C**, Same as panel A for networks trained to store *p* = 40 memories (*α* = 0.4). **D**, Same as panel A for networks trained to store *p* = 50 memories (*α* = 0.5). Empty panels indicate parameter combinations for which no solution was found. **E**, Reliability (cf. Fig. 2K) of the biophysical model versus the reliability of the corresponding binary model (from panels A–D). **F**, Reliability of the biophysical model (same as panel E) as a function of memory-pattern stability in the binary network. A minimum binary-network stability is required for successful recall in the biophysical model. **G**, Reliability of the biophysical model (same as panel E) as a function of memory load, *α* = *p/N*, for different activity levels.

**FIG. S6.**
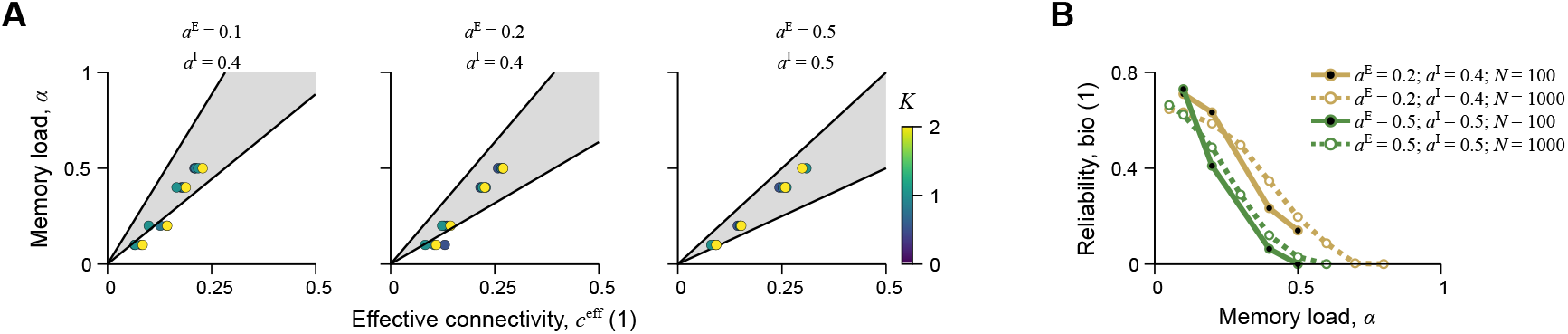
Approximate memory capacity for different parameters. **A**, Memory load, *α* = *p/N*, in binary-network simulations (circles) as a function of effective connectivity, *c*_eff_ = (*N*^E^*c*^E^ + *N*^I^*c*^I^)*/N*, shown for different activity levels (*a*^E^, *a*^I^). Circle colour denotes the modPLR stability parameter, *K*, used in Eqs. 52 and 53. Network size is *N* = 100. The shaded region indicates the interval between the lower and upper zero-stability capacity references obtained from the analytical approximation in the Supplementary Text. These references are first defined in terms of the separate excitatory and inhibitory connectivities, (*c*^E^, *c*^I^), and then projected onto the one-dimensional plotting variable *c*_eff_. **B**, Reliability of recall in the biophysical model as a function of memory load, shown for different activity levels and network sizes. Reliability decreases with increasing memory load, but this decrease is less abrupt in larger networks.

**FIG. S7.**
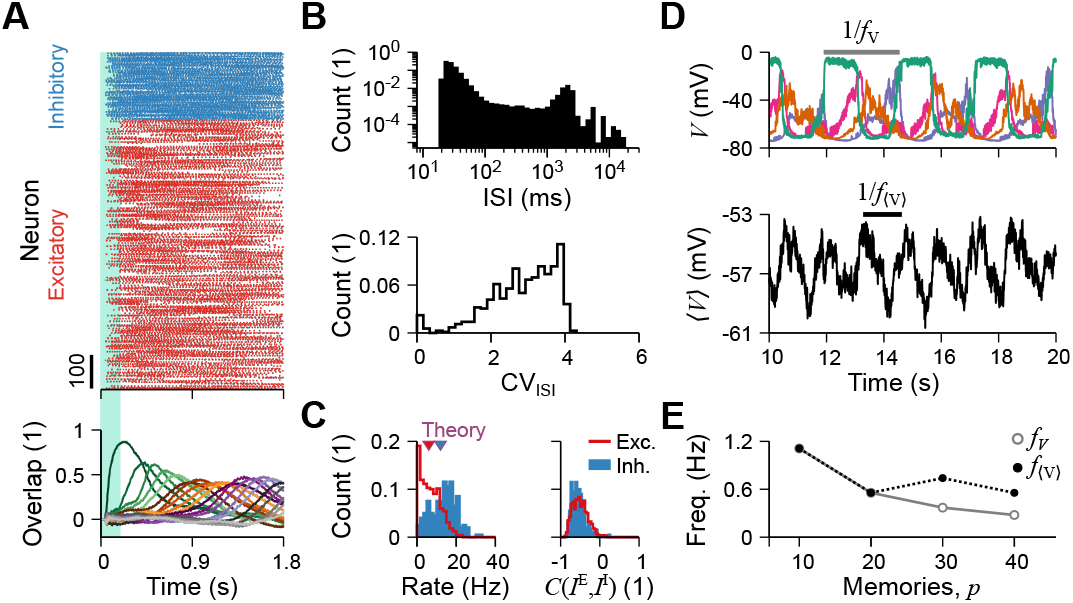
Autonomous sequential recall in biophysical memory networks. **A**, Spike times of excitatory (red) and inhibitory (blue) neurons (top) and memory overlap (bottom) as a function of time. The shaded area indicates the period of external stimulation. **B**, Histograms of interspike intervals (ISIs; top) and of the coefficient of variation of the ISIs (CV_ISI_; bottom). **C**, Histograms of average neuronal firing rates (left) and of the Pearson correlation between excitatory and inhibitory currents, *C*(*I*^E^, *I*^I^) (right). **D**, Temporal evolution of the dendritic membrane potential for four selected neurons (top) and of the average membrane potential across all excitatory neurons (bottom). The indicated quantities correspond to the oscillation periods, 1*/ f*_*V*_ and 1*/ f* ⟨_*V*_⟩, for the individual and average membrane potentials, respectively. **E**, Oscillation frequencies of the individual and average membrane potentials as a function of the number of stored memories, *p*.

**FIG. S8.**
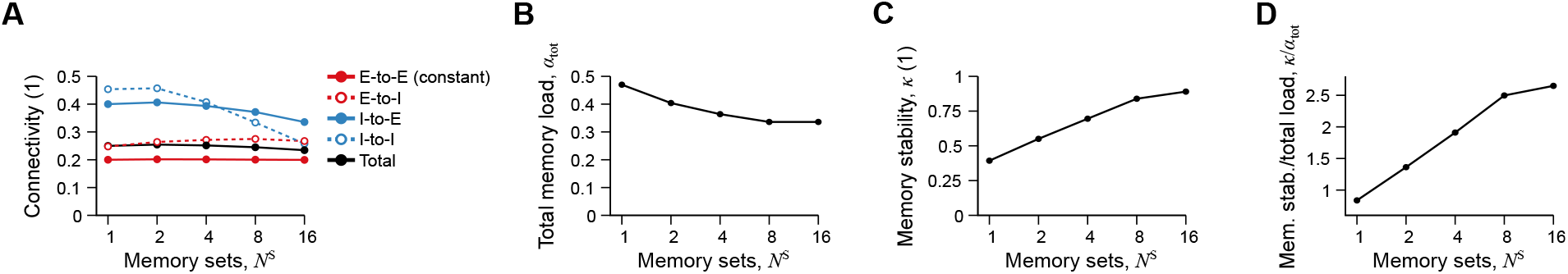
Subdividing memories into sets improves memory stability at fixed overall connection density. **A**, Connectivity of excitatory-to-excitatory (E-to-E), excitatory-to-inhibitory (E-to-I), inhibitory-to-excitatory (I-to-E), inhibitory-to-inhibitory (I-to-I), and total connectivity as a function of the number of memory sets. The modified perceptron learning rule was used to keep overall E-to-E connectivity constant. **B**, Total memory load of the network, *α*_tot_, as a function of the number of memory sets, using the same simulations as in panel A. The total memory load is the number of memories stored per set, *p*, multiplied by the number of memory sets, *N*^S^, and normalised by network size, *N*: *α*_tot_ = *pN*^S^*/N*. Increasing the number of memory sets leads to a small decrease in total memory load. **C**, Average stability per memory as a function of the number of memory sets, using the same simulations as in panel A. **D**, Average stability per memory divided by total memory load as a function of the number of memory sets, using the same simulations as in panel A.

**FIG. S9.**
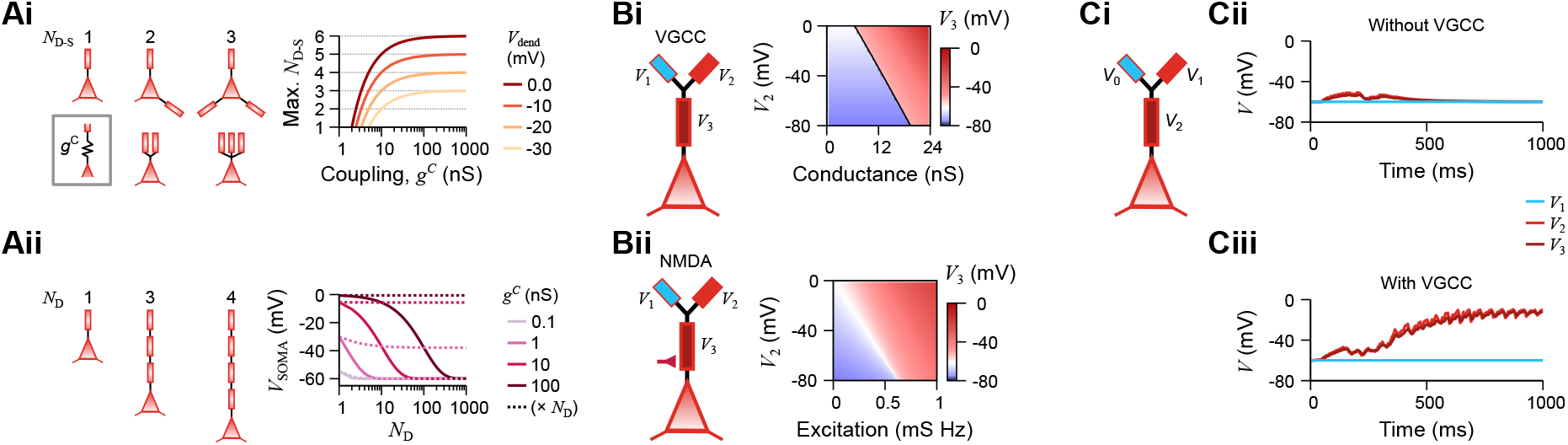
Active amplification in intermediate dendrites is required to store multiple memory sets. **A**, *Ai, left:* Schematics of *N*_D_−_S_ dendrites connected to the soma in parallel. Passive dendrite–soma coupling is denoted by *g*^C^. *Ai, right:* Maximum number of parallel dendrites, *N*_D_−_S_, for which the neuron still spikes when only one dendrite is depolarised, while all others remain hyperpolarised, as a function of passive dendrite–soma coupling, *g*^C^, for different dendritic depolarisation levels. *Aii, left:* Schematics of *N*_D_ dendrites connected to the soma in series. *Aii, right:* Steady-state somatic membrane potential as a function of the number of dendrites connected in series, *N*_D_, when the leaf branch is held at *V*_1_ = 0 mV, for different values of passive compartment–compartment coupling, *g*^C^ (coloured solid lines; see legend). Dashed lines show the corresponding somatic membrane potentials when the compartment–compartment coupling is multiplied by the number of dendrites. **B**, *Bi, left:* Schematic of a neuron with three dendritic branches. *Bi, right:* Steady-state membrane potential of the trunk dendritic branch (*V*_3_; colour) as a function of VGCC conductance (x-axis) and membrane potential of the depolarised dendrite (*V*_2_; y-axis), with *V*_1_ held at rest. The black line indicates *V* = −50 mV, the spiking threshold used here. *Bii*, Same as panel Bi, but for NMDA conductance on the trunk dendritic branch. The x-axis denotes NMDA excitation (cf. Fig. 1D). **C**, Temporal evolution of the membrane potentials of the three dendritic compartments highlighted in the schematics (Ci), without VGCC channels (only passive propagation; Cii) and with VGCC channels in the trunk dendritic branch (Ciii).

**FIG. S10.**
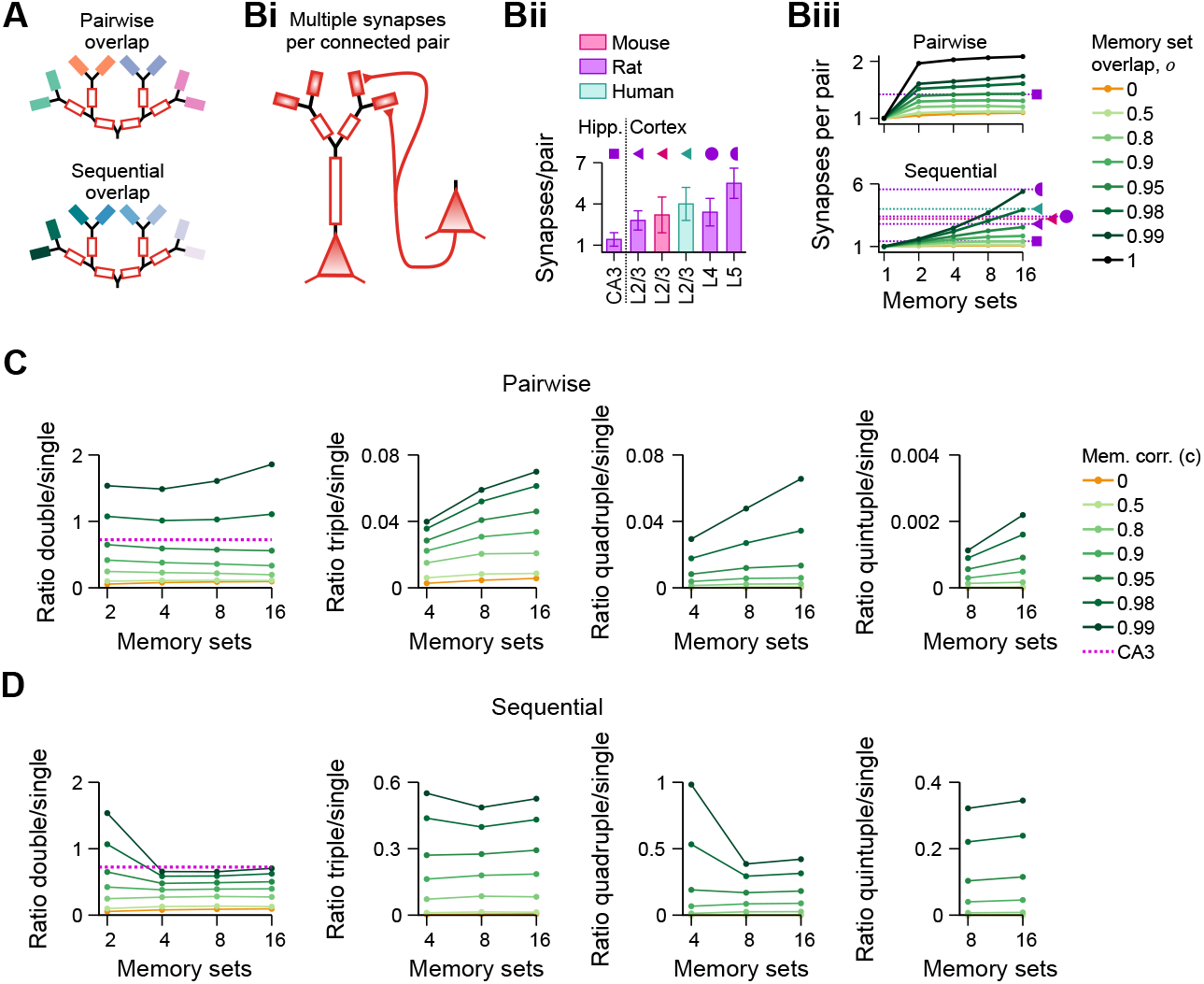
Overlap architecture predicts multi-synapse connectivity. **A**, Overlap architectures for dendritic memory sets. In the pairwise architecture (top), adjacent leaf dendrites share correlated memory patterns. In the sequential architecture (bottom), each memory set overlaps with the next one in the sequence. **B**, Multi-synapse connectivity. *Bi*, Schematic of a presynaptic neuron connecting to two postsynaptic dendrites. *Bii*, Number of synapses per connected pair of excitatory neurons in rat CA3^21^ and cortex: mouse and human L2/3^98^, rat L2/3^99^, rat L4^100^, and rat L5^101^. *Biii*, Predicted number of synapses per connected pair of excitatory neurons for different numbers of memory sets, overlap architectures, and overlap levels. Dashed horizontal lines denote the empirical reference values from panel Bii. **C**, Ratio of multiple to single synapses (double, triple, quadruple, quintuple) for different numbers of memory sets and overlap levels in the pairwise architecture (panel A, top). The dashed line indicates the CA3 reference value from ref. 21. **D**, Same as panel C, but for the sequential architecture (panel A, bottom). The dashed line indicates the CA3 reference value from ref. 21.

**FIG. S11.**
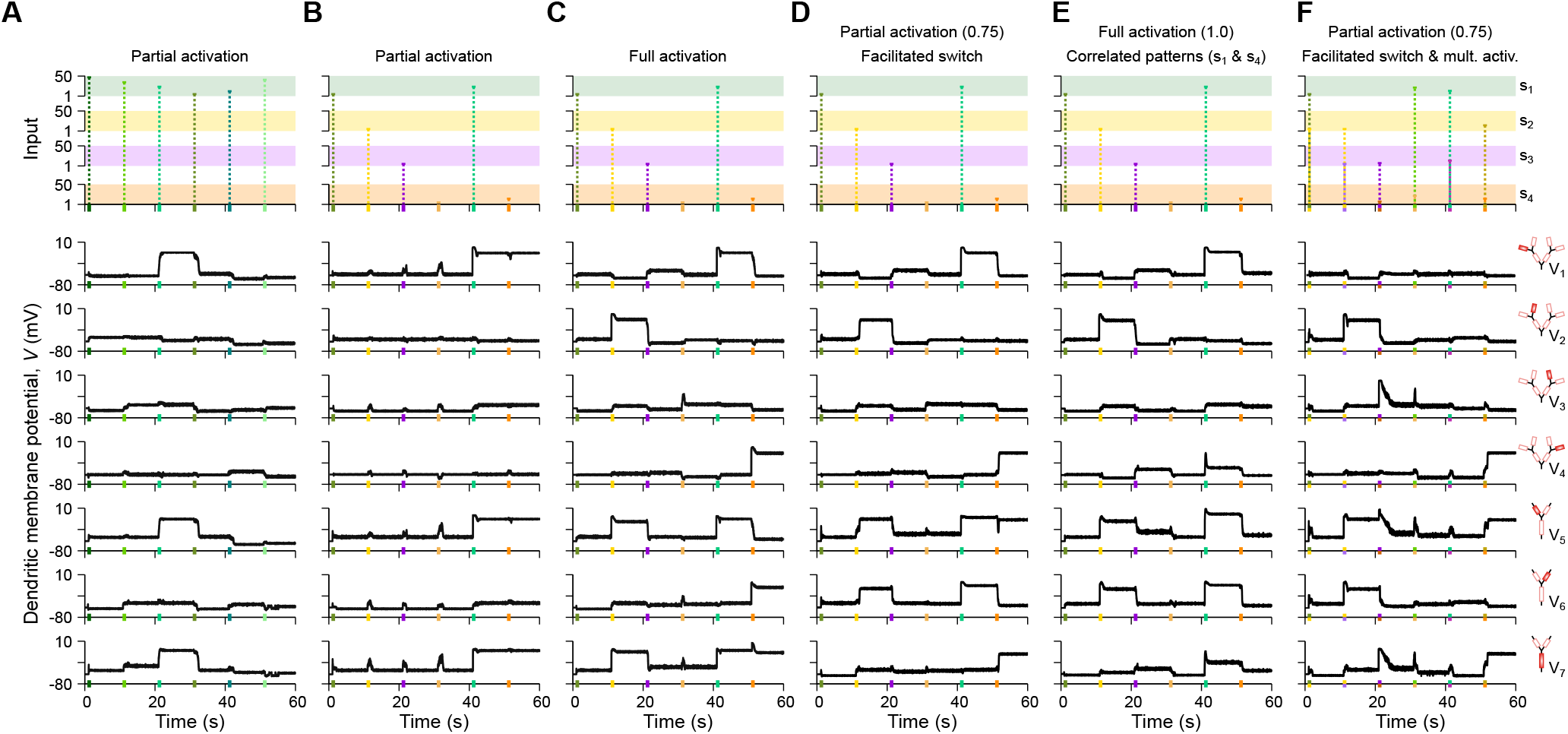
Dendritic membrane potentials of an example neuron in the autonomous system. **A**, Input to the memory network (top) and membrane potentials of the six dendrites of an example memory-network neuron (bottom), corresponding to Fig. 5C. **B**, Same as panel A, corresponding to Fig. 5D. **C**, Same as panel A, corresponding to Fig. 5E. **D**, Same as panel A, corresponding to Fig. 5F. **E**, Same as panel A, corresponding to Fig. 5G. **F**, Same as panel A, corresponding to Fig. 5H.

## Supplementary tables

**TABLE S1.** Simulation parameters for the neuron model.

| Parameter | Symbol | Value | Figs. |
| --- | --- | --- | --- |
| Membrane time constant | $\tau^M$ | 30 ms | 1–5; S1, S2, S3–S7, S9–S11 |
| Leak conductance per area | $\bar{g}^L$ | 10 nS/cm<sup>2</sup> | 1–3; S1, S2, S4, S9 |
| Membrane capacitance per area | $C$ | 300 pF/cm<sup>2</sup> | not explicitly used |
| Resting potential | $V^R$ | –60 mV | 1–5; S1, S2, S3–S7, S9–S11 |
| AMPA reversal potential | $E^A$ | 0 mV | 1–5; S1, S2, S3–S7, S9–S11 |
| NMDA reversal potential | $E^N$ | 0 mV | 1–5; S1, S2, S3–S7, S9–S11 |
| GABA<sub>A</sub> reversal potential | $E^{G_A}$ | –80 mV | 1–5; S1, S2, S3–S7, S9–S11 |
| GABA<sub>B</sub>/KiR reversal potential | $E^{K/G_B}$ | –90 mV | 1–5; S1, S2, S3–S7, S9–S11 |
| α5-GABA<sub>A</sub> reversal potential | $E^{G_\alpha}$ | –80 mV | 3–5; S11 |
| AMPA time constant | $\tau^A$ | 5 ms | 1–5; S1, S2, S3–S7, S9–S11 |
| NMDA time constant | $\tau^N$ | 150 ms | 1–5; S1, S2, S3–S7, S9–S11 |
| GABA<sub>A</sub> time constant | $\tau^{G_A}$ | 10 ms | 1–5; S1, S2, S3–S7, S9–S11 |
| GABA<sub>B</sub>/KiR time constant | $\tau^{K/G_B}$ | 150 ms | 1–5; S1, S2, S3–S7, S9–S11 |
| α5-GABA<sub>A</sub> time constant | $\tau^{G_\alpha}$ | 30 ms | 3–5; S11 |
| NMDA nonlinearity parameter 1 | $a^N$ | 0.15 | 1–5; S1, S2, S3–S7, S9–S11 |
| NMDA nonlinearity parameter 2 | $b^N$ | –12.5 mV | 1–5; S1, S2, S3–S7, S9–S11 |
| GABA<sub>B</sub>/KiR nonlinearity parameter 1 | $a^{K/G_B}$ | $e^{10}$ | 1–5; S1, S2, S3–S7, S9–S11 |
| GABA<sub>B</sub>/KiR nonlinearity parameter 2 | $b^{K/G_B}$ | 10 mV | 1–5; S1, S2, S3–S7, S9–S11 |
| α5-GABA<sub>A</sub> nonlinearity parameter 1 | $a^{G_\alpha}$ | $1.57 \times 10^{-7}$ | 3–5; S10 |
| α5-GABA<sub>A</sub> nonlinearity parameter 2 | $b^{G_\alpha}$ | –3 mV | 3–5; S11 |
| Ratio NMDA to AMPA | $R^{N/A}$ | 1 | 1; S1 |
|  |  | 0.5 | 2–5; S1, S2, S3–S7, S9–S11 |
| Ratio GABA<sub>B</sub>/KiR to GABA<sub>A</sub> | $R^{G_B/G_A}$ | 1 | 1; S1 |
|  |  | 0.5 | 2–5; S1, S2, S3–S7, S9–S11 |
| VGCC conductance | $\bar{g}^{Ca}$ | 60 | 5; S9, S11 |
| VGCC reversal potential | $E^{Ca}$ | 40 mV | 5; S9, S11 |
| VGCC nonlinearity parameter 1 | $a^{Ca}$ | 0.15 | 5; S9, S11 |
| VGCC nonlinearity parameter 2 | $b^{Ca}$ | –12.5 mV | 5; S9, S11 |
| Spiking threshold | $V^{TH}$ | –50 mV | 1–5; S1, S2, S3–S7, S9–S11 |
| Somatic reset potential | $V^{RT}$ | –60 mV | 1–5; S1, S2, S3–S7, S9–S11 |
| Somatic refractory period | $\tau^R$ | 5 ms | 1–5; S1, S2, S3–S7, S9–S11 |
| Coupling soma-dendrite | $\bar{g}^C$ | 0.5 | 2; S4, S9C |
|  |  | 1 | S7 |
|  |  | 1.5 | 3; S5, S6 |
|  |  | 0.41 | 5; S11 |
|  |  | 5 | S5, S6 |
| Coupling dendrite-dendrite | $\bar{g}_d^C$ | 50 | 5; S11 |
|  |  | 5 | S9 |
| Fluorescence signal time constant | $\tau^F$ | 150 ms | 4 |
| Fluorescence signal noise standard deviation | $\sigma^e$ | 0.001 | 4 |

**TABLE S2.**
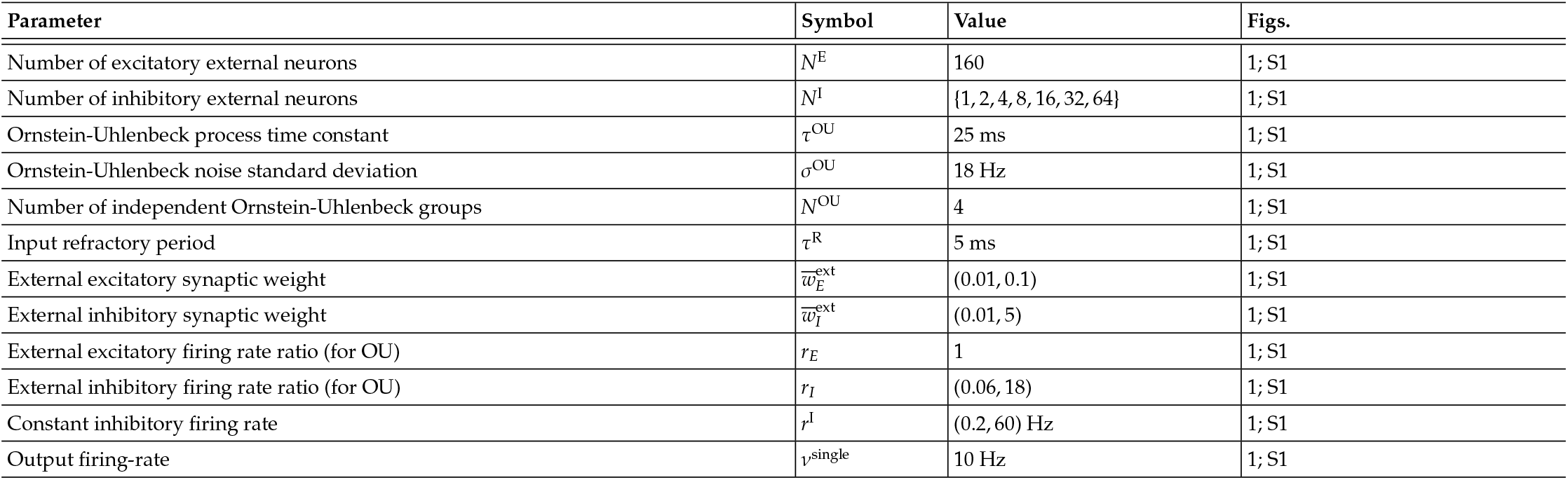
Simulation parameters for the single neuron.

**TABLE S3.**
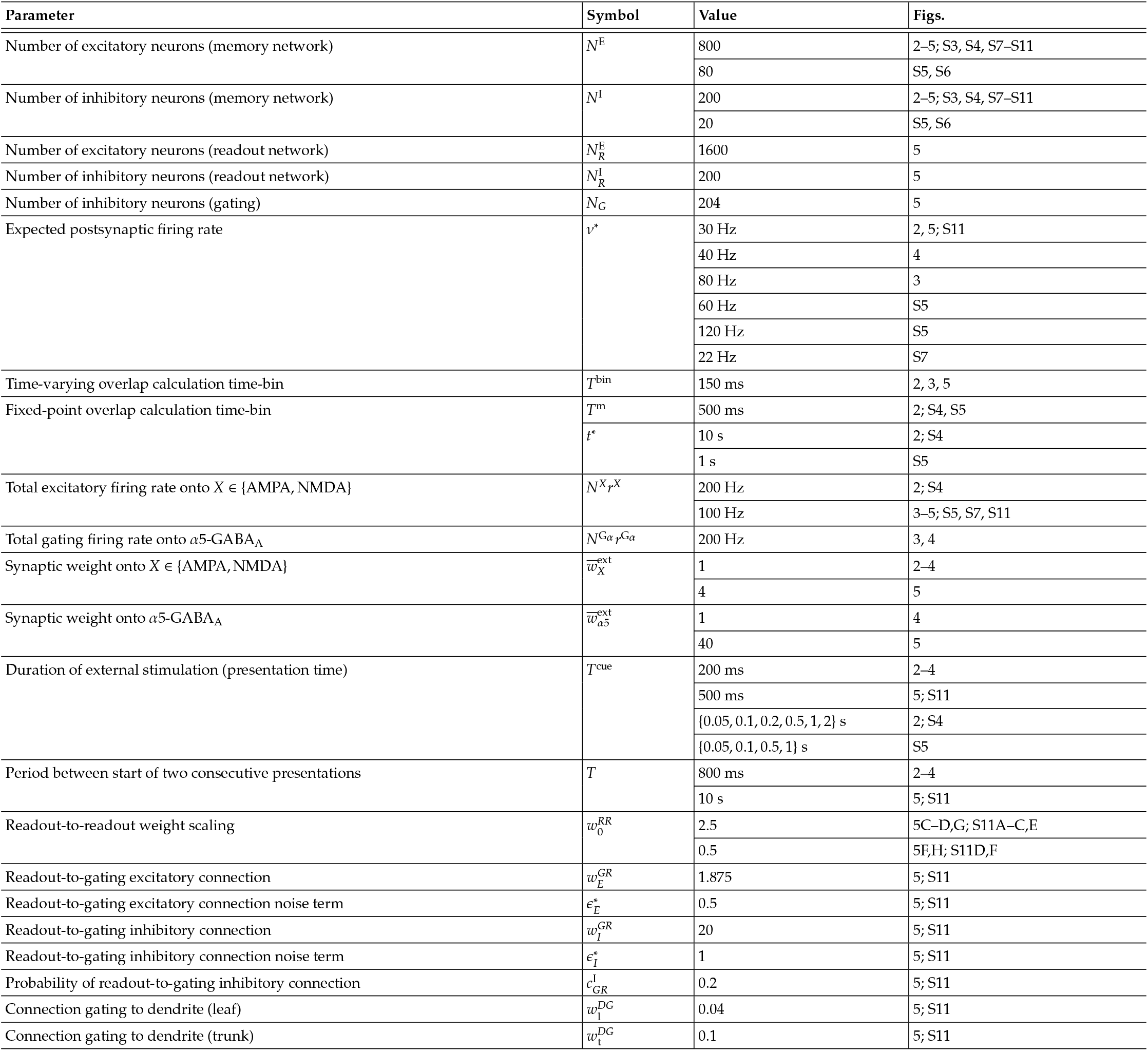
Simulation parameters for the recurrent network.

**TABLE S4.**
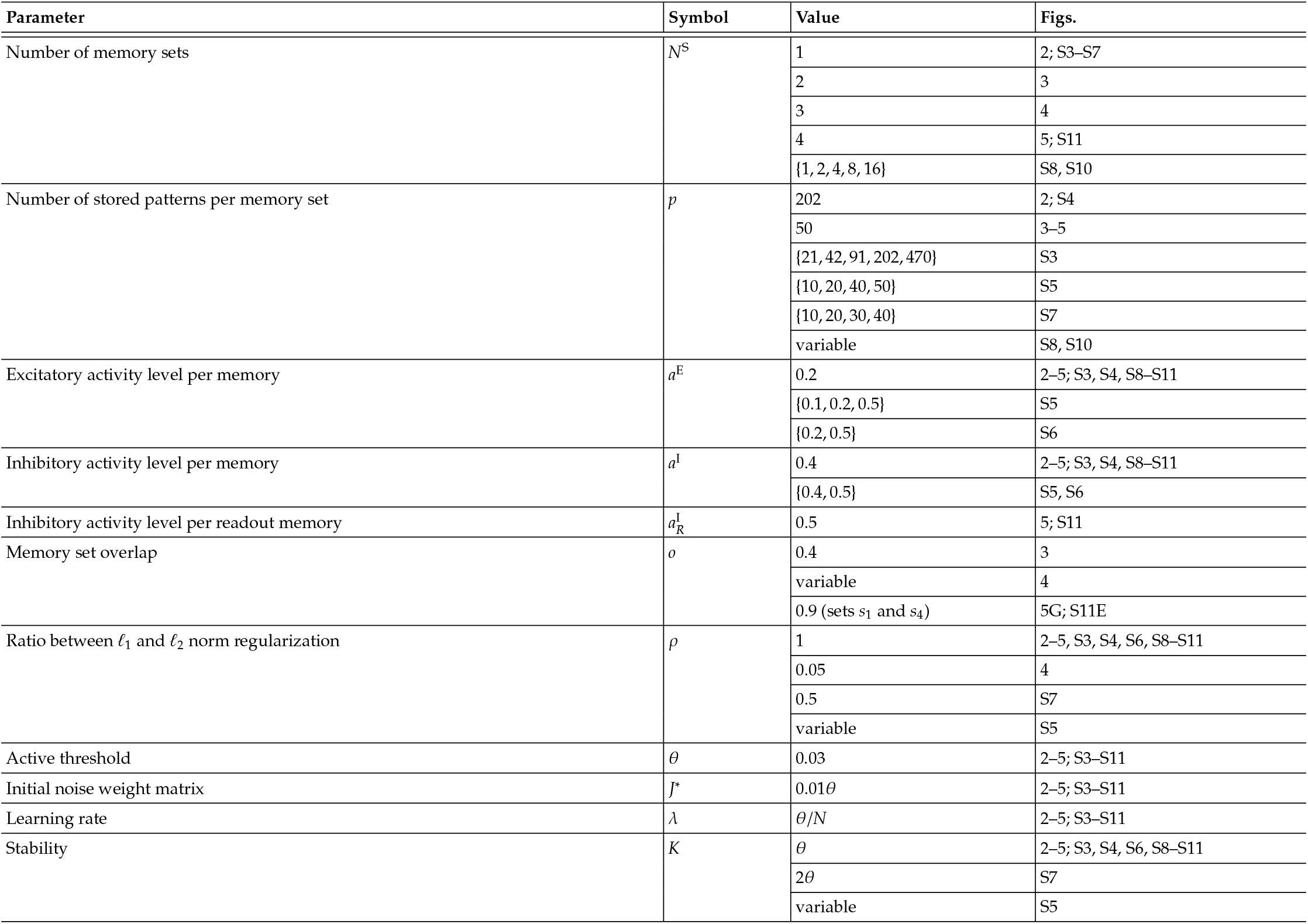
Simulation parameters for memories.

## Supplementary text

### Attractor landscape and pseudo-energy

The landscapes shown schematically in Fig. 3F are intended as an intuitive representation of the attractor structure induced by the effective connectivity, rather than as a rigorous Lyapunov function. In symmetric associative-memory networks, stored memories correspond to minima of a true energy landscape. In our framework, however, the connectivity does not need to be symmetric, so a global scalar energy function is not generally defined. We therefore use the term *pseudo-energy* only as a visual proxy for the effective attractor landscape: stable recalled memories correspond to basins of attraction, whereas inhibitory gating changes the effective connectivity and thereby switches which set of attractors is accessible at a given time.

### Additional analyses of dendritic integration

#### Maximum number of dendrites and minimum dendritic coupling for passive dendritic integration

In our simplified biophysical neuron model, a spike is generated when the somatic membrane potential crosses the spike threshold, *V*^TH^, from below. For this to occur, the current arriving from the immediately connected trunk dendrite must be sufficient to depolarise the soma above threshold. Different dendritic architectures are possible, including parallel and serial configurations, both of which attenuate depolarisation from leaf dendrites in the absence of active ion channels such as VGCCs.

#### Parallel dendrites connected directly to the soma

For multiple dendrites connected directly to the soma in parallel, at least one dendritic compartment must be depolarised, noting that the soma itself does not receive excitatory synaptic input. Following our inhibitory-gating scheme, if *N*^D−S^ dendritic compartments are connected directly to the soma in parallel, only one is depolarised at a given time, while the remaining *N*^D−S^ − 1 dendritic compartments are hyperpolarised. The somatic membrane potential then obeys

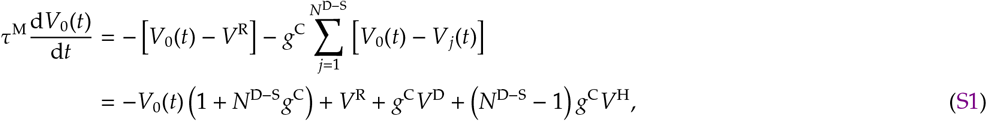

where *V*^D^ and *V*^H^ are the membrane potentials of the depolarised and hyperpolarised dendritic compartments, respectively. To determine the maximum number of dendrites that can be connected in parallel while still allowing the soma to emit at least one spike, we set Eq. S1 to zero and *V*_0_ = *V*^TH^, yielding

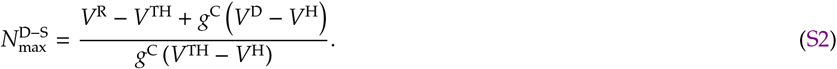

The maximum number of dendritic compartments connected directly to the soma depends on the dendrite–soma coupling. In the limit of infinite coupling, the upper bound becomes

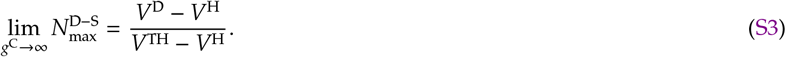

Similarly, the minimum dendrite–soma coupling required for the soma to spike when only one dendritic compartment is depolarised is

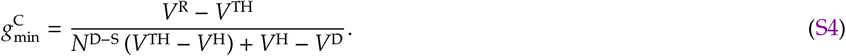

#### Serially connected dendrites

For multiple dendrites connected unidirectionally in series, the membrane potential of dendrite *i* obeys

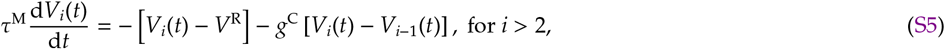

where *V*_*i*_(*t*) is the membrane potential of dendrite *i*, which receives passive current from dendrite *I* − 1. We are interested in the steady state and therefore take the leaf dendrite, indexed by *i* = 1, to be fully depolarised, 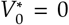 mV. Under this condition, Eq. S5 has the solution

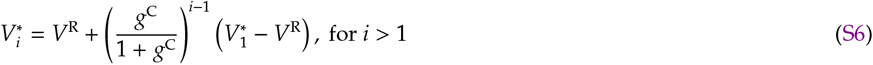

With *N*^D^ dendrites, the steady-state somatic membrane potential becomes

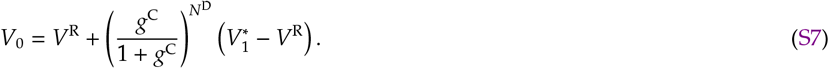

### Additional analyses of the binary model

#### Recurrent memory-encoding weights for sequences

To store autonomous recall of memory sequences, we modify the conditions (Eqs. 52 and 53) as

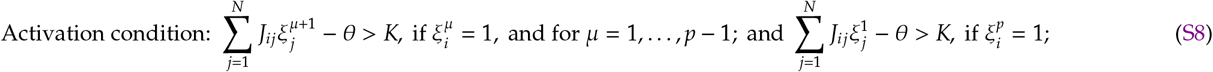

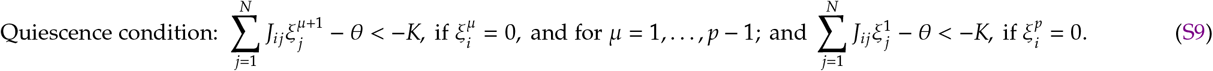

All other modPLR steps remain the same as described in the Methods section.

#### Increase in memory stability and sparsity

Our modified Perceptron Learning Rule (modPLR) increases memory stability while reducing connectivity. These two objectives are promoted by decreasing the *ℓ*_2_ and *ℓ*_1_ norms of the input vectors of all units, respectively. We therefore define the loss function

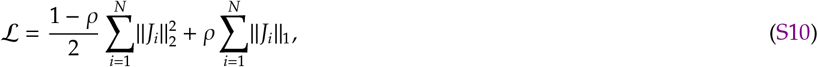

where

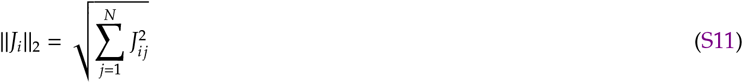

and

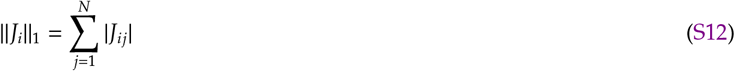

are the *ℓ*_2_ and *ℓ*_1_ norms of the input connectivity vector of unit *i*, respectively. The gradient of this loss function is

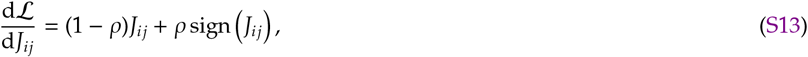

which yields the shrinkage update

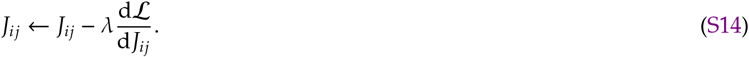

In the implementation used here, this shrinkage step is applied after convergence of the modPLR update rules and initiates a new optimisation iteration.

#### Multiple synaptic contacts

Networks encoding multiple memory sets across complex dendritic trees can exhibit distinct overlap architectures, such as *pairwise* or *sequential* organisation. In the pairwise case, overlap is restricted to specific partners, for example leaf dendrites sharing the same branching point, while all other pairs remain independent (Fig. S10A, top). In the sequential case, memory sets are arranged in a linear chain such that each set overlaps with the next (Fig. S10A, bottom). Although membrane potential and calcium fluorescence depend primarily on the degree of overlap in both architectures (cf. Fig. 4F), the corresponding structural connectivity differs. In particular, the prevalence of multiple synaptic contacts between connected neuron pairs depends on the overlap architecture (Fig. S10Bi), enabling comparison with experimental measurements (Fig. S10Bii)^2198–101^. Accordingly, the number of synapses per connected pair depends on the degree of overlap, the number of dendritic leaves, and the overall overlap architecture (Fig. S10Biii). Under these assumptions, pairwise overlap reproduces the average number of synaptic contacts observed in CA3^21^, whereas sequential overlap is additionally consistent with reported cortical connectivity values^98–101^ (Fig. S10Biii). Because pairwise architectures link only specific pairs of dendritic leaves, three or more contacts per connected pair remain rare even as additional memory sets are added (Fig. S10C). By contrast, sequential architectures can generate a substantial probability of three or more contacts per connected pair (Fig. S10D).

### Approximate memory capacity

#### Definitions

Here we construct a zero-stability approximation for the memory capacity of the binary network defined in the Methods section. The approximation is based on the sign-constrained perceptron theory of Chapeton *et al*. ^16^ . For clarity, we repeat below the definitions needed for this construction.

For our calculations, we consider one postsynaptic unit—which represents a randomly selected excitatory or inhibitory unit from the network—and denote its incoming weight vector by **J**, where *J*_*j*_ is the weight from presynaptic unit *j*. The first *N*^E^ presynaptic units are excitatory and the remaining *N*^I^ are inhibitory, with

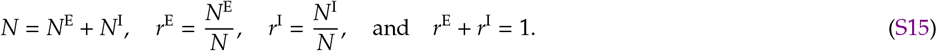

The network stores *p* binary memory patterns, denoted by 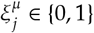 for *µ* = 1, …, *p*. Their activity levels are given by Eq. 44,

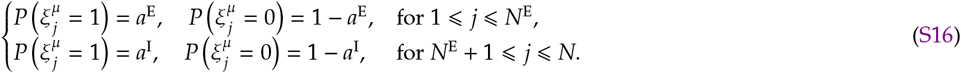

In the corresponding single-unit classification problem, the output activity is denoted by *a*°. For an excitatory postsynaptic unit, *a*° = *a*^E^, whereas for an inhibitory postsynaptic unit, *a*° = *a*^I^. We characterize structural sparsity by the excitatory and inhibitory connection densities, *c*^E^ and *c*^I^, respectively. We define the structural effective connectivity as

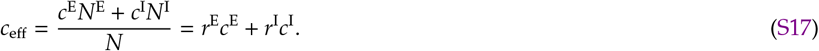

Thus, *Nc*_eff_ is the expected total number of incoming synapses per postsynaptic unit.

#### Zero-stability single-population slope

We focus on the zero-stability case (*K* = 0 in Eqs. 52 and 53) and quantify memory load by *α* = *p/N*. Rather than repeating the full statistical-mechanics derivation of Chapeton *et al*. ^16^, we start directly from their final zero-stability system of equations. In that framework, the output activity enters through the scalar equation

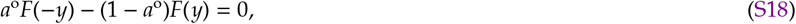

where

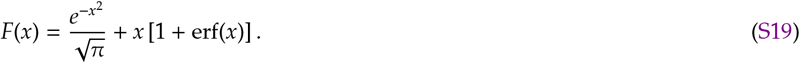

For each activity level *a*°, Eq. S18 defines a unique value of *y*. Excitatory and inhibitory populations enter through the additional condition

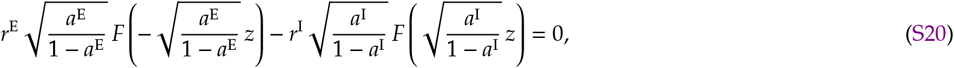

which defines the corresponding value of *z* for the chosen population statistics (*r*^E^, *r*^I^, *a*^E^, *a*^I^). The associated zero-stability critical capacity is then^16^

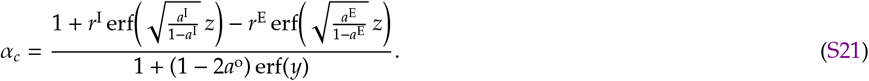

Note that connectivity is not a parameter in the calculations of Chapeton *et al*. ^16^, but a resulting property of a network with parameters *a*^E^, *a*^I^, *r*^E^, *r*^I^, and a given memory stability. This constrains the space of solutions. For example, for *a*^E^ = *a*^I^ = 0.5, and *r*^E^ = 0.8 and *r*^I^ = 0.2, excitatory and inhibitory connectivity densities are 0 ⩽ *c*^E^ ⩽ 0.5 and 0.5 ⩽ *c*^I^ ⩽ 1. Our modPLR allows for sparse connectivity for both excitatory and inhibitory connections, which requires additional assumptions expanded below.

#### Approximate capacity

To isolate the dependence of the zero-stability capacity on the output activity, we now simplify the numerator of Eq. S21. Because erf(*x*) ∈ [−1, 1] and *r*^E^ + *r*^I^ = 1, the numerator is bounded above by

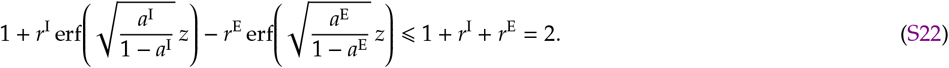

We denote this maximal value by

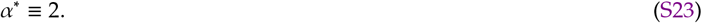

Using this bound, Eq. S21 implies the output-activity upper envelope

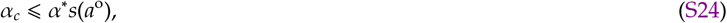

where we define

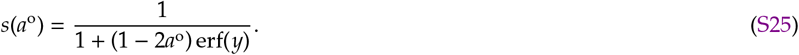

We use *s*(*a*°) as an activity-dependent capacity factor for a single postsynaptic population. For the two populations in the network, we therefore define

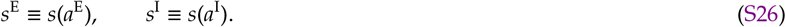

This step isolates the dependence of the zero-stability branch on the output activity, but discards the explicit *z*-dependent asymmetry between excitatory and inhibitory populations.

We now reintroduce the two-population structure of the network by approximating the capacity factor as the sum of separate excitatory and inhibitory contributions. Specifically, we assume that the retained excitatory and inhibitory synapses contribute additively, with weights determined by their relative abundances in the network, *r*^E^ and *r*^I^, and by the corresponding activity-dependent capacity factors, *s*^E^ and *s*^I^. This gives the aggregate two-population approximation

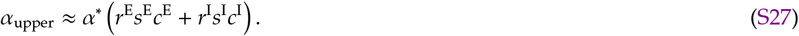

For the purpose of the one-dimensional representation in Fig. S6, we further approximate this expression by replacing the connectivity-dependent slope with the weighted average

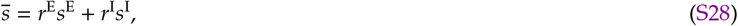

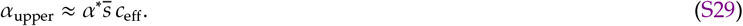

As a lower reference, we consider instead the case in which the retained non-zero synapses are placed at random. Within the present zero-stability approximation, we assume that the activity-dependent capacity factors remain the same as in the upper reference, but that random placement reduces the overall capacity by a factor of two relative to the optimistic prefactor *α*^∗^. This choice recovers the sign-constrained unbiased case when *a*^E^ = *a*^I^ = 0.5, for which *s*^E^ = *s*^I^ = 1 and the lower reference reduces to *c*_eff_. We therefore write

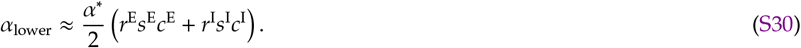

Using the same one-dimensional projection as above, this becomes

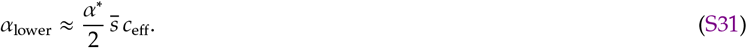

Thus, after projection onto *c*_eff_, the one-dimensional upper slope is a connectivity-weighted average of the excitatory and inhibitory single-population slopes. A stricter bottleneck approximation would instead replace this weighted average by the more restrictive population, for example through min(*s*^E^, *s*^I^). In the present simulations, however, the modified perceptron learning rule changes not only the capacity factors *s*^E^ and *s*^I^, but also the learned connectivities *c*^E^ and *c*^I^ themselves. As a result, differences in coding affect both the predicted slope and the position of simulation points along the *c*_eff_ axis. For this reason, we use the weighted approximation above when comparing simulation points in the (*c*_eff_, *α*) plane. The underlying approximation nevertheless remains two-dimensional in (*c*^E^, *c*^I^), and the projection onto *c*_eff_ is used only for visualization.

